# Intestinal barrier function in the naked mole-rat: an emergent model for gastrointestinal insights

**DOI:** 10.1101/2024.01.17.576063

**Authors:** Javier Aguilera-Lizarraga, Anne Ritoux, David C. Bulmer, Ewan St. John Smith

## Abstract

The intestinal barrier plays a crucial role in homeostasis, both by facilitating absorption of nutrients and fluids, and providing a tight shield to prevent the invasion by either pathogen or commensal microorganisms. Intestinal barrier malfunction is associated with systemic inflammation, oxidative stress, and decreased insulin sensitivity, which may lead to the dysregulation of other tissues. Therefore, a deeper understanding of physiological aspects related to an enhanced barrier function is of significant scientific and clinical relevance. The naked mole-rat has many unusual biological features, including attenuated colonic neuron sensitivity to acid and bradykinin, and resistance to chemical-induced intestinal damage. However, insight into their intestinal barrier physiology is scarce. Here, we observed notable macroscopic and microscopic differences in intestinal tissue structure between naked mole-rats and mice. Moreover, naked mole-rats showed increased number of larger goblet cells and elevated mucus content. In measuring gut permeability, naked mole-rats showed reduced permeability compared to mice, measured as transepithelial electrical resistance, especially in ileum. Furthermore, intestinal ion secretion induced by serotonin, bradykinin, histamine, and capsaicin was significantly reduced in naked mole-rats compared to mice, despite the expression of receptors for all these agonists. In addition, naked mole-rats exhibited reduced pro-secretory responses to the non-selective adenylate cyclase activator forskolin. Collectively, these findings indicate that naked mole-rats possess a robust and hard-to-penetrate gastrointestinal barrier, that is resistant to environmental and endogenous irritants. Naked mole-rats may therefore provide valuable insights into the physiology of the intestinal barrier and set the stage for the development of innovative and effective therapies.

**Graphical abstract:** 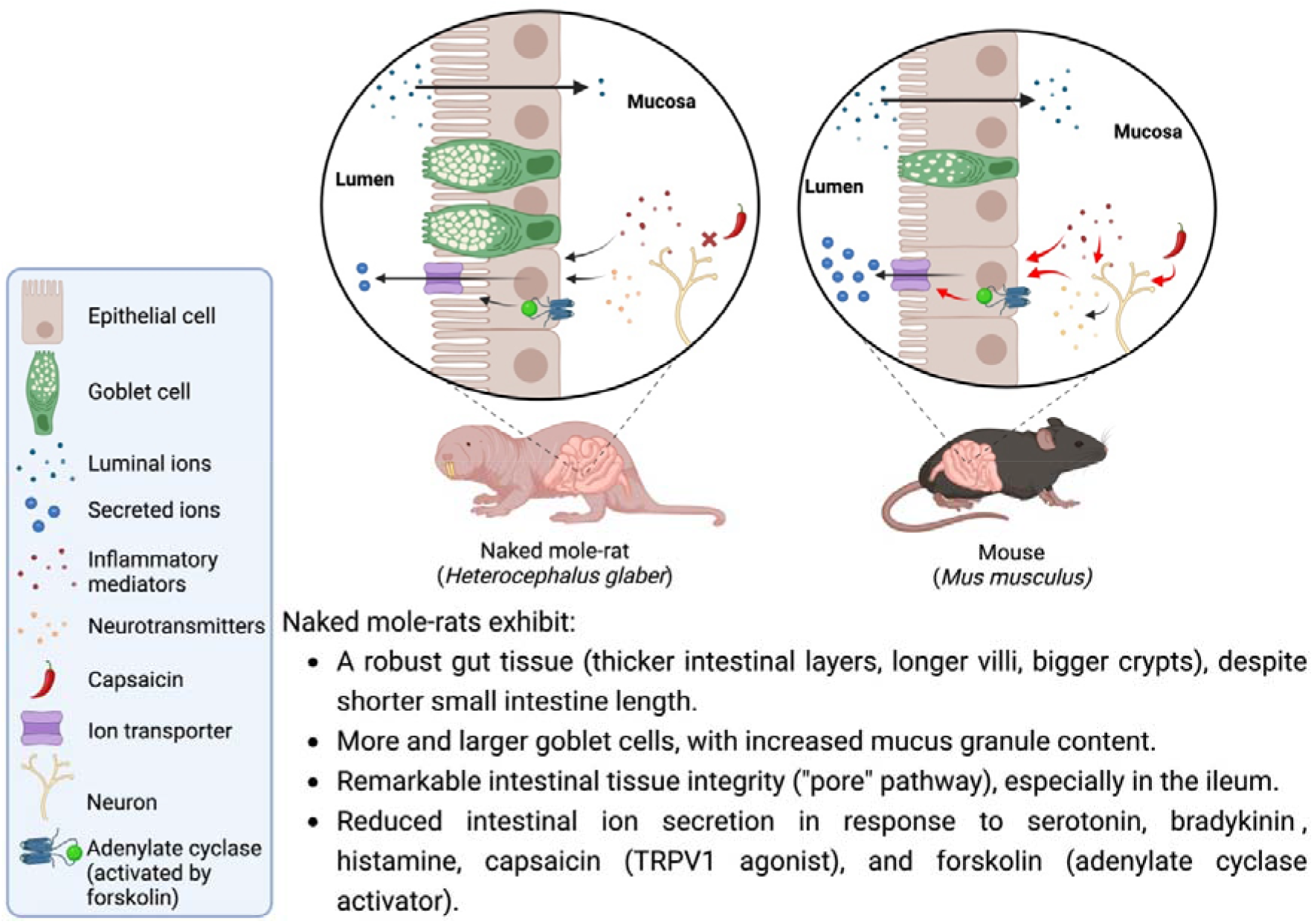

## 1. Introduction

Digestive pathologies stand out as the most common type of disease worldwide (*1*). Importantly, they result in a significant reduction in the quality of life for patients and represent a multi-billion healthcare burden globally (*2–5*). Intestinal barrier dysfunction is a common feature of most gastrointestinal diseases, including coeliac disease, inflammatory bowel disease, metabolic dysfunction-associated liver disease and disorders of gut-brain interaction (*6*). Moreover, numerous pathologies unrelated to the gastrointestinal tract, such as respiratory diseases (*7*), skin conditions (*8*) and neurodegenerative diseases (*9*), have also been associated with impaired intestinal barrier function. The common denominator of intestinal barrier perturbation led to the “leaky gut” hypothesis (*10*), which states that enhanced intestinal permeability, caused by endogenous or exogenous factors, may increase the passage of food antigens, commensal microorganisms and/or pathogens into the underlying tissue and the systemic circulation, leading to aberrant immune activation described in different disease conditions. Although increased permeability may not necessarily be deleterious (*11*), and whether intestinal barrier dysregulation is the cause or consequence in most disorders is unknown, it is broadly accepted that adequate functioning of the intestinal barrier is required to maintain gut and inter-organ homeostasis. Consequently, identifying mechanisms leading to enhanced intestinal barrier function is highly relevant for both basic and clinical sciences.

In addition to the use of human intestinal mucosal biopsies (*12*), many models have been employed to investigate the (patho)physiology of the intestinal barrier. Among these, human gut-derived cell lines (*13*), such as Caco-2 cell monolayers (*14*), and animal models, primarily rodents, but also porcine, equine and avian models (*15*), have been most commonly utilised. Furthermore, human organoids have recently been implemented to model intestinal epithelial barrier function (*16*, *17*). Despite important advances in our understanding of intestinal barrier function, there has been little to no reduction in the global incidence and prevalence of gastrointestinal conditions over the last 30+ years (*1*). There is thus an imperative need for further research, including the development of novel models, to enhance understanding of intestinal barrier function.

In recent years, naked mole-rats (NMR, *Heterocephalus glaber*) have attracted considerable attention from biomedical scientists as a non-traditional animal model due to their highly unusual biological features (*18*). Notably, these animals show an outstanding tolerance to environmental stressors associated with human diseases. Indeed, these eusocial rodents, which have an unusually long lifespan given their 40-80 g size (>35 years) (*19*), have been used for investigating ‘healthy’ aging (*20*), cancer resistance (*21*), and protection against neurodegeneration (*22*). However, the gastrointestinal tract of these animals remains poorly characterised. Of interest, a recent study revealed that the intestinal barrier of NMRs is resistant to tissue damage induced by dextran sodium sulphate (DSS) (*23*), a widely used colitogenic chemical for modelling inflammatory bowel disease. Although the authors employed a modified version of the traditional DSS protocol (*24*), NMRs displayed no signs of histological damage, even when exposed to 8.75% DSS, albeit for only 3-days. Although several factors may be involved in the increased resistance to DSS observed in these animals, such as a thicker mucus layer (*23*), differences in the gut microbiota (*25*), a distinct immune system (*26*), etc., this observation suggests that the gut physiology of NMRs may possess characteristics of potential biological interest. We therefore hypothesised that NMRs would display enhanced intestinal function, that is, improved permeability regulation and resistance to intestinal irritants. To answer this question, we explored the macroscopic and microscopic features of the NMR gut, intestinal permeability, changes in mucosal ion secretion, and expression of intestinal markers of biological interest. Investigating the intestinal barrier function in NMRs may enable a better understanding of the underlying mechanisms that contribute to better mucosal protection.

## 2. Material and Methods

### 2.1. Animals

All animal experiments were performed in accordance with the Animals (Scientific Procedures) Act 1986 Amendment Regulations 2012 under Project Licenses granted to E.St.J.S. (P7EBFC1B1 and PP5814995) by the Home Office and approved by the University of Cambridge Animal Welfare Ethical Review Body. Experiments were performed using a mixture of male and female C57BL6/J mice (10-15 weeks old; IQR 10-12) and a mixture of male and female, non-breeder, NMRs (23-145 weeks; IQR 48-54). Mice were purchased from Envigo and housed conventionally with nesting material and a red plastic shelter in a temperature-controlled room at 21 °C, with a 12-hour light/dark cycle and access to food and water *ad libitum*. NMRs were bred in-house and maintained in an interconnected network of cages in a humidified (∼55%) temperature-controlled room at 28-32 °C with red lighting (08:00−16:00). NMRs had access to food *ad libitum*. In addition, a heat cable provided extra warmth under two to three cages/colony. NMRs used in this study came from five different colonies. Animals were humanely killed by CO_2_ exposure followed by cervical dislocation (mice) or decapitation (NMRs).

### 2.2. Intestinal tissue processing for histology and histomorphometric assessment

Intestinal biopsies (ileum and colon) were fixed and embedded using conventional techniques. In brief, samples were incubated, after dissection, in 4% paraformaldehyde (PFA) at 4 °C overnight. Next, samples were washed in PBS, transferred to sucrose (0.3 g · ml^−1^) and stored at 4 °C overnight. Finally, samples were embedded in Shandon M-1 Embedding Matrix (Thermo Fisher Scientific), snap-frozen and stored at −80°C until processing. Cryostat (Leica) sections (12 μm) were collated across Superfrost Plus slides (Thermo Fisher Scientific) and stained with haematoxylin and eosin (H&E) using conventional techniques (*27*, *28*). Villi length and width, crypts length and width, and thickness of the different layers (submucosa and muscularis) were measured in ileum and colon samples (Supplementary Figure 1a, b) using ImageJ software. For analyses of villi and crypts, only intestinal units that were overtly complete and full-sized, and showed no signs of bending or mechanical damage, were chosen in each sample. A total of 5-10 units of each kind were carefully selected, and their average calculated.

### 2.3. Alcian blue/periodic acid-Schiff (AB/PAS) staining

For goblet cell identification and quantification of their mucus content, 12-μm tissue sections were subjected to Alcian Blue and Periodic Acid Schiff’s (AB/PAS) staining. In brief, sections were hydrated in double distilled water and then stained with Alcian Blue solution (Sigma, 1% w/v Alcian Blue in 3% v/v aqueous acetic acid) for 15 minutes. Next, sections were rinsed in double distilled water before being treated for 5 minutes with 0.5% periodic acid (Sigma). Thereafter, tissue sections were washed again in double distilled water and stained for 10 minutes in Schiff’s reagent (Sigma). Finally, sections were washed for 5 minutes in tap water, mounted, and imaged using a NanoZoomer S360 (Hamamatsu).

For analyses, images were loaded into the QuPath software (*29*). In each image, regions of interest (ROI) for crypts and villi were manually annotated. Within these, the cell detection tool was used to segment individual cells using a manually selected threshold. To ensure that the segmentation correctly circled cells, ROIs were visually inspected. After colour deconvolution, measurements for average pixel intensity across each cell ROI and cell area were analysed.

### 2.4. Ussing chambers

Post-mortem, for both species, the gastrointestinal tract (from pyloric sphincter to anus) was carefully removed, cleaned of faecal pellets, and maintained in ice-cold Krebs solution. For experiments where transepithelial electrical resistance (*TEER*) was measured, full-thickness segments of ileum and colon were mounted in Ussing chambers without removal of the muscularis propria or the serosa. For experiments where short-circuit current (*I_sc_*) was measured, ileum samples were pinned in Sylgard plates with the lamina propria facing down and muscularis externa and serosal layers gently peeled off with forceps to prepare lamina propria-submucosal plexus intestinal strips. Multichannel DVC 1000 Voltage/Current clamp apparatus (World Precision Instruments) was used to measure changes in intestinal tissue function. Each channel was connected to a DVC-3 preamplifier (World Precision Instruments), connected to two current and two voltage electrodes filled with 1.5% (w/v) agar (Millipore) dissolved in 3M KCl. The voltage and fluid resistance were zeroed to eliminate electrical bias and the voltage clamp circuitry was set to standby mode and the intestinal preparation was mounted in the chamber with fresh superfusate. After calibration, intestinal tissues were then mounted in Ussing chambers and the mucosal and serosal compartments were bathed independently with 10 mL of Krebs-Ringer bicarbonate buffer (pH 7.4) containing (in mM): 118 NaCl; 4.7 KCl; 1.2 CaCl_2_; 1.2 MgSO_4_; 1.2 NaH_2_PO_4_; 25 NaHCO_3_). Glucose (10 mM) was present in the medium that bathed the serosal surface of the tissue and mannitol (10 mM) was substituted for glucose in the medium that bathed the mucosal surface. Media were oxygenated (95% O_2_, 5% CO_2_) throughout the experiment via a gas-uplift recirculation system and maintained at 37°C by water-jacketed reservoirs. Although NMRs are cold blooded and maintained at 28-32 °C, to minimise the number of variables impacting our results, all experiments were conducted with media maintained at 37 °C. For mouse tissue, Ussing chambers with an exposed area of 0.071 cm^2^ were used, while an exposed area of 0.196 cm^2^ was used for NMR tissue.

#### Transepithelial electrical resistance (TEER) experiments

The transepithelial electrical resistance was calculated according to Ohm’s law (resistance = voltage/current, normalised by the exposed window area; TEER = R_tissue_(Ω) · Area (mm^2^)) from the voltage deflections induced by bipolar current pulses of 50 μA every 50 s with a duration of 500 ms. After 30 min of tissue stabilisation, TEER was measured over a period of 2 h.

#### Short-circuit current (I_sc_) experiments

After mounting, tissue was allowed to equilibrate for 15-20 min. Thereafter, preparations were short-circuited (voltage-clamped at zero potential) and the short-circuit current (*I_sc_*) required to maintain the 0 mV potential was monitored over time. After short-circuiting, intestinal tissue was allowed to equilibrate for 10-15 min to allow bioelectric stabilisation (baseline). All *I_sc_* responses were normalised to μA · cm^-2^ and changes in *I_sc_* (Δ*I_sc_*) were recorded after application of compounds to the serosal side, except forskolin which was added to both the apical and the serosal sides.

### 2.5. Chemicals

The following drugs were used for short-circuit current (*I_sc_*) experiments: capsaicin (Sigma-Aldrich), bradykinin (Tocris), 5-hydroxytryptamine (5-HT, serotonin) (Sigma-Aldrich), histamine (Sigma-Aldrich) and forskolin (Tocris). Capsaicin (1 mM) was dissolved in 100% ethanol, forskolin was dissolved in dimethyl sulfoxide (DMSO, 10 mM), and bradykinin (1 mM), 5-HT (10 mM) and histamine (10 mM) were dissolved in double distilled water.

### 2.6. RNA extraction from intestinal tissue and cDNA preparation

Following dissection and cleaning, intestinal samples (ileum and colon) were stored in RNAprotect Tissue Reagent (Qiagen) at 4°C until processing. Next, total RNA was isolated using the RNeasy Mini Kit (Qiagen) following the manufacturer’s instructions and including a DNase digestion step. Subsequently, cDNA was transcribed using qScript cDNA Synthesis Kit (Quantabio) following the manufacturer’s instructions.

### 2.7. Real-time quantitative PCR (RT-qPCR)

Reactions were prepared with FastStart Sybr Green Master (Merk) following the manufacturer’s instructions. In brief, 10 μL reaction mix per well, containing cDNA (20 ng) oligonucleotides (200 nM) and UltraPure distilled water (DNase/RNase free; Invitrogen), were loaded onto 96-well plates. Thermocycling was performed using a StepOnePlus^TM^ Real-Time PCR System (Thermo Fisher). A melt curve was performed at the end of thermocycling to assess the specificity of primers. Single peaks were obtained for all primer pairings.

To determine primer efficiency, a standard curve was generated for each pair of primers at cDNA concentrations of 50, 5, 0.5, 0.05 and 0.005 ng/µL and plotting C_t_ against log concentration. Efficiency was calculated by the following equation: Efficiency = 10^−(1/slope)^-1 (*30*, *31*). Efficiency of reaction values between 90 and 110% were considered acceptable.

Gene expression was normalised to the endogenous reference gene *Tbp* for both species. This gene encodes the TATA-binding protein. *Tbp* was chosen as it has been proposed as an ideal reference gene for the normalisation of gene expression in intestinal samples of mice and other animals (*32–36*). Relative gene expression was calculated as 2^−ΔΔCt^ (*37*). The primer pairs used are described in Supplementary Table 1.

### 2.8. Measurement of Substance P from intestinal explants

Post-mortem, the small intestine and colon were carefully removed, cleaned of faecal pellets and maintained in ice-cold Krebs solution. Next, 30-50 mg tissue sections (terminal ileum and distal colon) were dissected and transferred into 24-well plates containing 500 μL per well of L-15 Medium (GlutaMAX) medium (Gibco). Explants from both species were incubated for 5 min at 32 °C (5% CO_2_, 3%O_2_). Thereafter, culture supernatant was collected and stored at -80°C until use. Substance P was measured using an enzyme-linked immunosorbent assay (Substance P ELISA Kit, Cayman Chemicals) according to the manufacturer’s instructions. Substance P levels were normalised to protein content upon quantification using the Bradford Assay (Thermo Fisher) following the manufacturer’s instructions.

### 2.9. Statistical analyses

Tissue was taken from non-manipulated healthy animals being used for other experiments, thus meeting the Reduction aim of the 3Rs. While there were no upfront power calculations to determine the sample size, we included sample sizes commonly used for the types of assays used in this study (*38*, *39*).

Normal (Gaussian) distribution was determined for all datasets using the Shapiro–Wilk normality test. Non-parametric tests were used for datasets where ≥1 group did not pass the normality test (assuming α = 0.05). Otherwise, parametric tests were used. The type of statistical test and the sample sizes for each experiment are provided in figure legends. Data were plotted and statistical analyses performed with Prism (GraphPad Software version 10.2.1). A *p*-value <0.05 was considered statistically significant. Finally, effect sizes were calculated post *hoc* using the statistical software G*Power (Version 3.1.9.6).

For histological assessment of mouse and NMR samples, 2-4 sections per animal, separated by at least 150 μm, were randomly captured, and analysed by an experimenter in a blinded manner. The experimenter was unblinded for statistical analysis.

## 3. Results

### 3.1. Macroscopic features of the naked mole-rat gastrointestinal tract

NMRs are small rodents whose body mass is significantly larger compared to mice (data from this study: mouse 21.05 ± 1.71 g vs. NMR 47.07 ± 12.03 g, *p* < 0.0001, Figure 1a), yet substantially smaller than typically used laboratory adult rats (Sprague-Dawley, ∼300-400 g; Wistar rats, ∼250-500 g) (*40*, *41*) or guinea pigs (Hartley, ∼625-1,200 g) (*42*). Despite the larger size of NMRs compared to mice, NMRs had a noticeably shorter gut length, measured from the pyloric sphincter to the anus (mouse 40.09 ± 2.35 cm vs. NMR 22.17 ± 1.94 cm, *p* < 0.0001; Figure 1b, c). Further analyses revealed that these differences were driven by a shorter small intestine in NMR than that of mice (small intestine: mouse 32.69 ± 1.92 cm vs. NMR 14.95 ± 1.88 cm, *p* < 0.0001; colon: mouse 7.40 ± 0.65 cm vs. NMR 7.28 ± 0.59 cm, *p* = 0.6425; Figure 1d, e), in keeping with previous studies (*23*, *43*). Of interest, while the ratio of the small intestine/colon length in mice was comparable to that in humans (*44–46*), this was reduced by ∼2-fold in NMRs (human 3.956 ± 1.36 vs. mouse 4.52 ± 0.99 vs. NMR 2.07 ± 1.20, *p* < 0.0001; Figure 1f). The lengths of the NMR small intestine and colon were comparable between male and female (small intestine: male 15.50 ± 6.60 cm vs. female 15.05 ± 2.50 cm, *p* = 0.5626; colon: male 7.10 ± 2.00 cm vs. female 7.55 ± 0.20 cm, *p* = 0.2265; Supplementary Figure 1c, d). Given the considerable age range of the NMRs used in our study (minimum age: 23 weeks old; maximum age: 145 weeks old) and their attainment of adulthood at ∼1 year old, we analysed the relationship between gut length and both age and weight. The colon length was not associated with the age or weight of the NMR (age vs. length: r = 0.0581, p = 0.8248, weight vs. length: r = 0.0245, p = 0.9339; Supplementary Figure 1e, f). However, we did find statistically significant correlations between the small intestine length and the age and weight of NMRs (age vs. length: r = 0.7293, p = 0.0012, weight vs. length: r = 0.6258, p = 0.0282; Supplementary Figure 1g, h). These findings parallel human anatomical data, that is, the length of the colon in young/adolescent individuals is similar to that in adults (*47*), while the small intestine only reaches its maximum length in adulthood (*48*). Overall, these results show gross anatomical differences in the gut of NMRs compared to humans and mice, especially regarding the short length of the small intestine.

**Figure 1.**
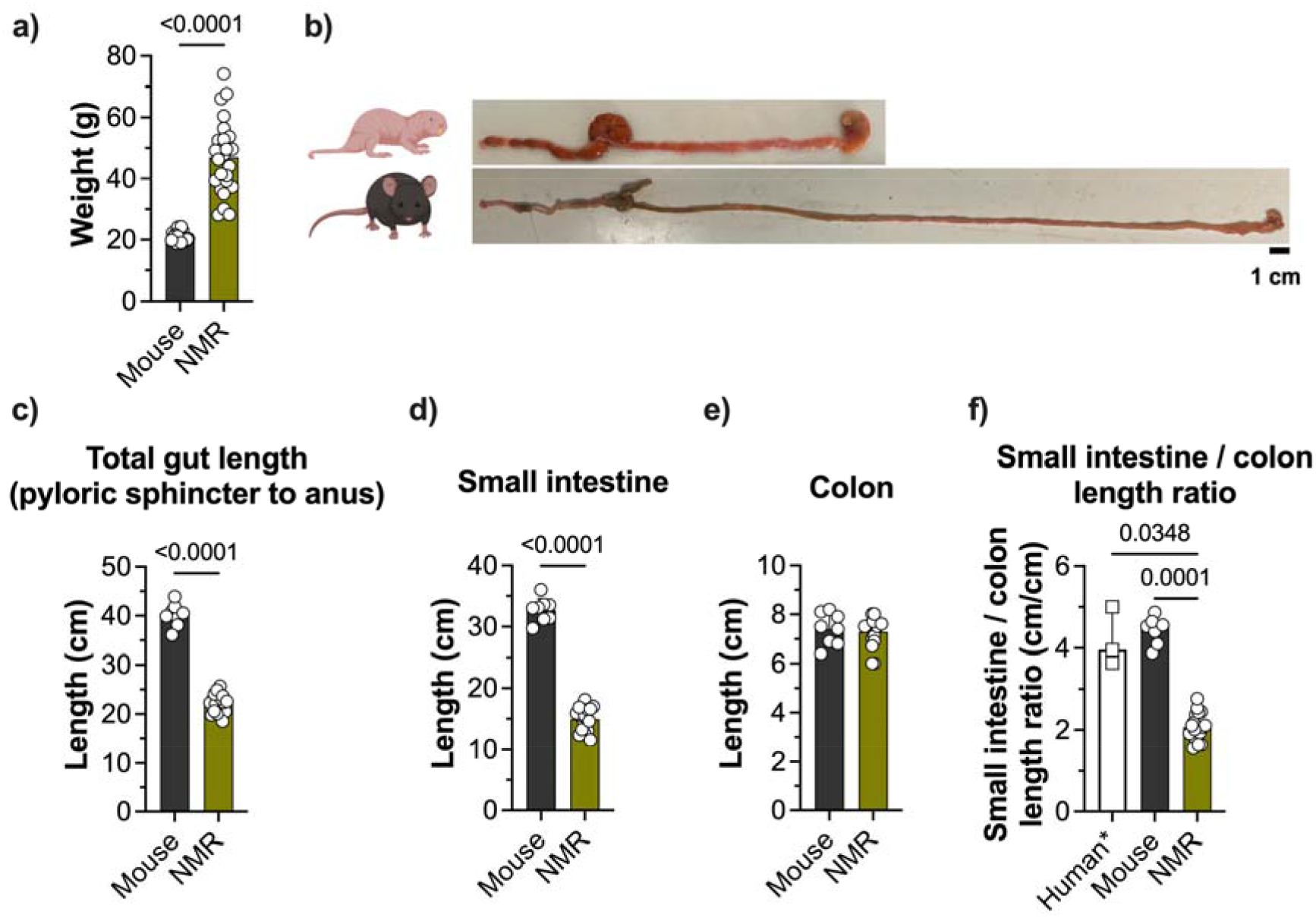
Anatomical features of gastrointestinal tracts of NMRs compared to mice. Weight of mice (n = 20) compared to NMR (n = 25) (**a**). **b**, Representative pictures of the gastrointestinal tract (from stomach to anus) of NMR (upper panel) and mouse (lower panel). Gastrointestinal length measured from the pyloric sphincter to the anus (**c**), small intestine (**d**) and colon (**e**) in mice (n= 8) and NMRs (n = 17). **f)** Length ratio of the small intestine compared to the colon in humans (data extracted from (*44–46*)), mice and NMR. Human* data reflect median of the means provided by 3 studies that analysed 100, 100 and 200 individuals, respectively. *P* values are shown in plots. **c**, **d** and **e**, unpaired t-test (data shown as mean ± SD); **f**, Kruskal-Wallis test followed by Dunn’s multiple comparisons test (data shown as median ± IQR). The NMR and mouse illustrations in **a** were created with BioRender.com).

### 3.2. Intestinal histology in naked mole-rats

Next, we explored the microscopic architecture of the small intestine (ileum) and colon in NMRs, comparing them to those of mouse tissue (Figure 2a-d). To this end, we measured the histological properties of the intestinal villi and crypts, as well as the thickness of different layers. The ileum of NMRs showed thicker muscularis (mouse 37.56 ± 8.24 μm vs. NMR 60.45 ± 23.43 μm, *p* = 0.0055; Figure 2e) and submucosal layers compared to mice (mouse 18.90 ± 2.02 μm vs. NMR 40.46 ± 11.22 μm, *p* < 0.0001; Figure 2f). Furthermore, NMR ileal villi were substantially longer (mouse 344.4 ± 180.8 μm vs. NMR 801.0 ± 394.3 μm, *p* < 0.0001; Figure 2g) and wider (mouse 90.95 ± 10.00 μm vs. NMR 135.3 ± 36.63 μm, *p* = 0.0008; Figure 2h) compared to those of mice, and NMR ileal crypts were significantly deeper (mouse -135.60 ± 66.81 μm vs. NMR -157.10 ± 47.49 μm, *p* = 0.0039; Figure 2i) and wider (mouse 35.72 ± 6.45 μm vs. NMR 57.86 ± 10.39 μm, *p* < 0.0001; Figure 2j). In the colon, the muscularis was comparable in both species (mouse 128.2 ± 78.31 μm vs. NMR 139.3 ± 83.82 μm, *p* = 0.2188; Figure 2k), but the colonic submucosa was thicker in NMRs (mouse 63.13 ± 10.18 μm vs. NMR 80.27 ± 10.26 μm, *p* = 0.0012; Figure 2l). Finally, colonic crypts were similarly deep in both species (mouse -219.0 ± 32.57 μm vs. NMR -198.0 ± 12.64 μm, *p* = 0.0837; Figure 2m), but those of NMRs were significantly wider compared to mice (mouse 34.07 ± 3.35 μm vs. NMR 56.79 ± 7.24 μm, *p* < 0.0001; Figure 2n).

**Figure 2.**
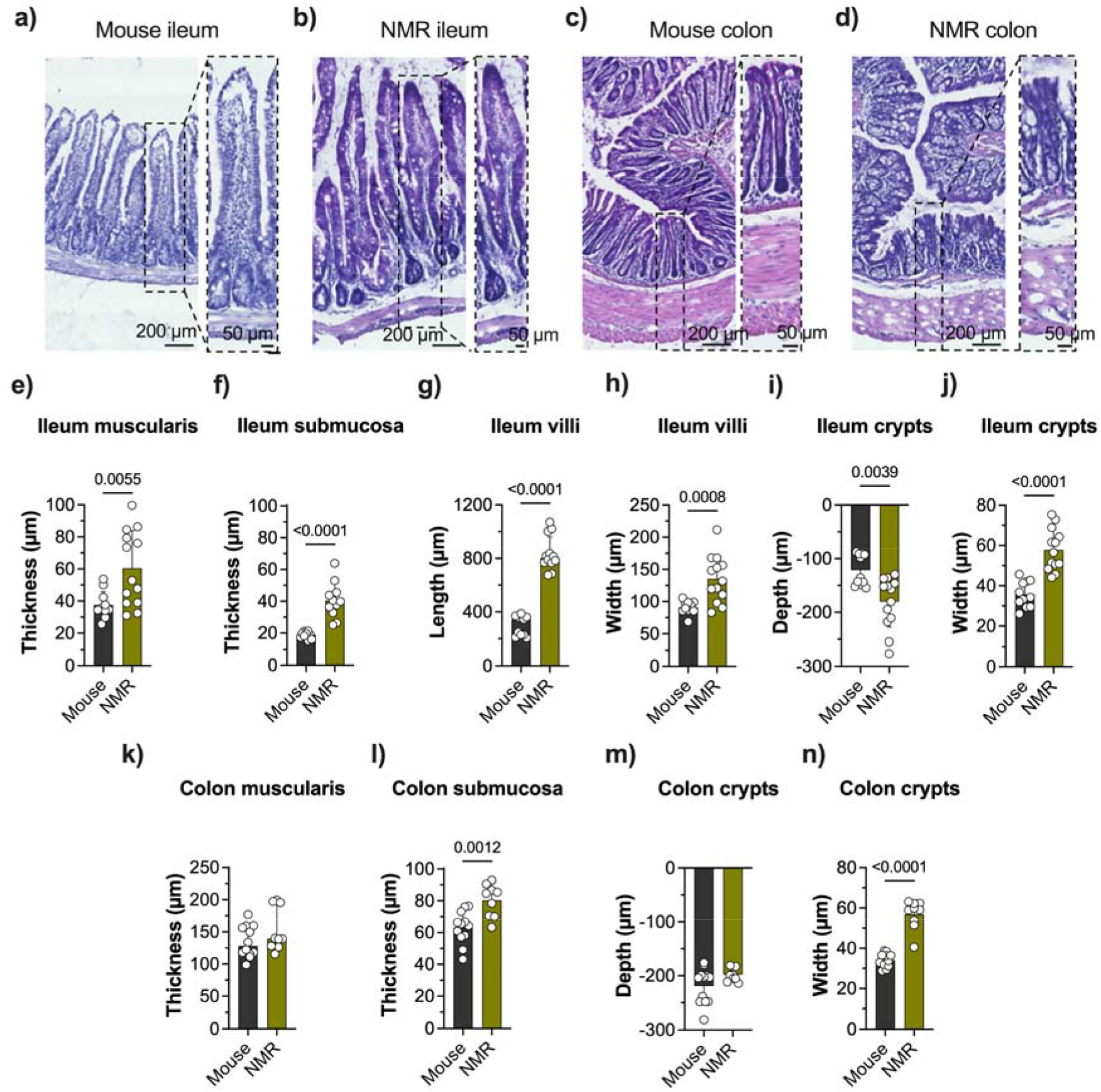
Intestinal histology in NMRs. Representative images of H&E staining sections from mouse (**a**) and NMR (**b**) ileum, and mouse (**c**) and NMR (**d**) colon. Ileum muscularis (**e**) and submucosal (**f**) layers thickness in mouse (n = 11, 2-3 sections/animal, 4 mice) and NMR (n = 13, 3-4 sections/animal, 4 NMR). **g**, average villi height (5-10 villi/sample) in mice (n = 11, 2-3 sections/animal, 4 mice) and NMRs (n = 12). **h**, average villi width (5-10 villi/sample) in mice (n = 11, 2-3 sections/animal, 4 mice) and NMRs (n = 13, 3-4 sections/animal, 4 NMR). **i**, average crypt depth (5-10 crypts/sample) in mice (n = 11, 2-3 sections/animal, 4 mice) and NMRs (n = 13, 3-4 sections/animal, 4 NMR). **j**, average crypt width (5-10 crypts/sample) in mice (n = 11) and NMRs (n = 13). Colon muscularis (**k**) and submucosal (**l**) layers thickness in mouse (n = 12, 3 sections/animal, 4 mice) and NMR (n = 9, 3 sections/animal, 3 NMRs). **m**, average crypt depth (5-10 crypts/sample) in mice (n = 12, 3 sections/animal, 4 mice) and NMRs (n = 9, 3 sections/animal, 3 NMRs). **n**, average crypt width (5-10 crypts/sample) in mice (n = 12, 3 sections/animal, 4 mice) and NMRs (n = 9, 3 sections/animal, 3 NMRs). *P* values are shown in plots. **e**, **f**, **h**, **j**, **l**, **m** and **n**, unpaired t-test (data shown as mean ± SD); **g**, **i** and **k**, Mann-Whitney test (data shown as median ± IQR).

We next used Alcian Blue/Periodic Acid-Schiff (AB/PAS) staining to identify goblet cells and quantify mucus granule content in small intestine and colon samples. Ileum samples (Figure 3a, b) of NMRs showed similar numbers of AB^+^ mucin-producing goblet cells in the villi compared to mice (mouse 80.07 ± 23.25 cells/villus vs. NMR 93.07 ± 20.41 cells/villus, *p* = 0.2254; Figure 3c), and slightly higher numbers in the crypts (mouse 28.66 ± 7.97 cells/crypt vs. NMR 38.98 ± 8.86 cells/crypt, *p* = 0.0194; Figure 3c). However, the size of the goblet cells present in the NMR ileum was significantly larger than that of mouse ileum goblet cells, both in the villi and crypts (mouse villi 33.91 ± 3.42 μm^2^ vs. NMR villi 41.79 ± 7.62 μm^2^, *p* = 0.012; mouse crypts 29.25 ± 3.71 μm^2^ vs. NMR crypts 46.53 ± 5.79 μm^2^, *p* < 0.0001; Figure 3d). In addition, the goblet cell-mucus granule content (measured as AB/PAS staining intensity per cell) in NMR ileum, in both villi- and crypts-residing cells, was significantly higher compared to that of mice (mouse villi 0.51 ± 0.04 a.u. vs. NMR villi 0.74 ± 0.10 a.u., *p* < 0.0001; mouse crypts 0.48 ± 0.06 a.u. vs. NMR crypts 0.89 ± 0.12 a.u., *p* < 0.0001; Figure 3e). In the colon (Figure 3f, g), the number of AB^+^ goblet cells was significantly elevated in the crypts of NMRs compared to in mice (mouse 14.07 ± 1.51 cells vs. NMR 23.97 ± 4.54 cells, *p* < 0.0001; Figure 3h). Furthermore, as in the ileum, the size of these goblet cells was significantly bigger in NMRs compared to mice (mouse crypts 65.98 ± 14.77 μm^2^ vs. NMR crypts 170.2 ± 21.43 μm^2^, *p* < 0.0001; Figure 3d). Finally, again, in keeping with the results obtained in the ileum, the mucus granule content in NMR colonic crypts was significantly larger than that of mouse colonic crypts (mouse 1.00 ± 0.05 a.u. vs. NMR 1.42 ± 0.19 a.u., *p* < 0.0001; Figure 3j). Interestingly, differences in the AB/PAS staining colouration between these two species may suggest that their mucin compositions differ, but further investigation is required to substantiate this.

**Figure 3.**
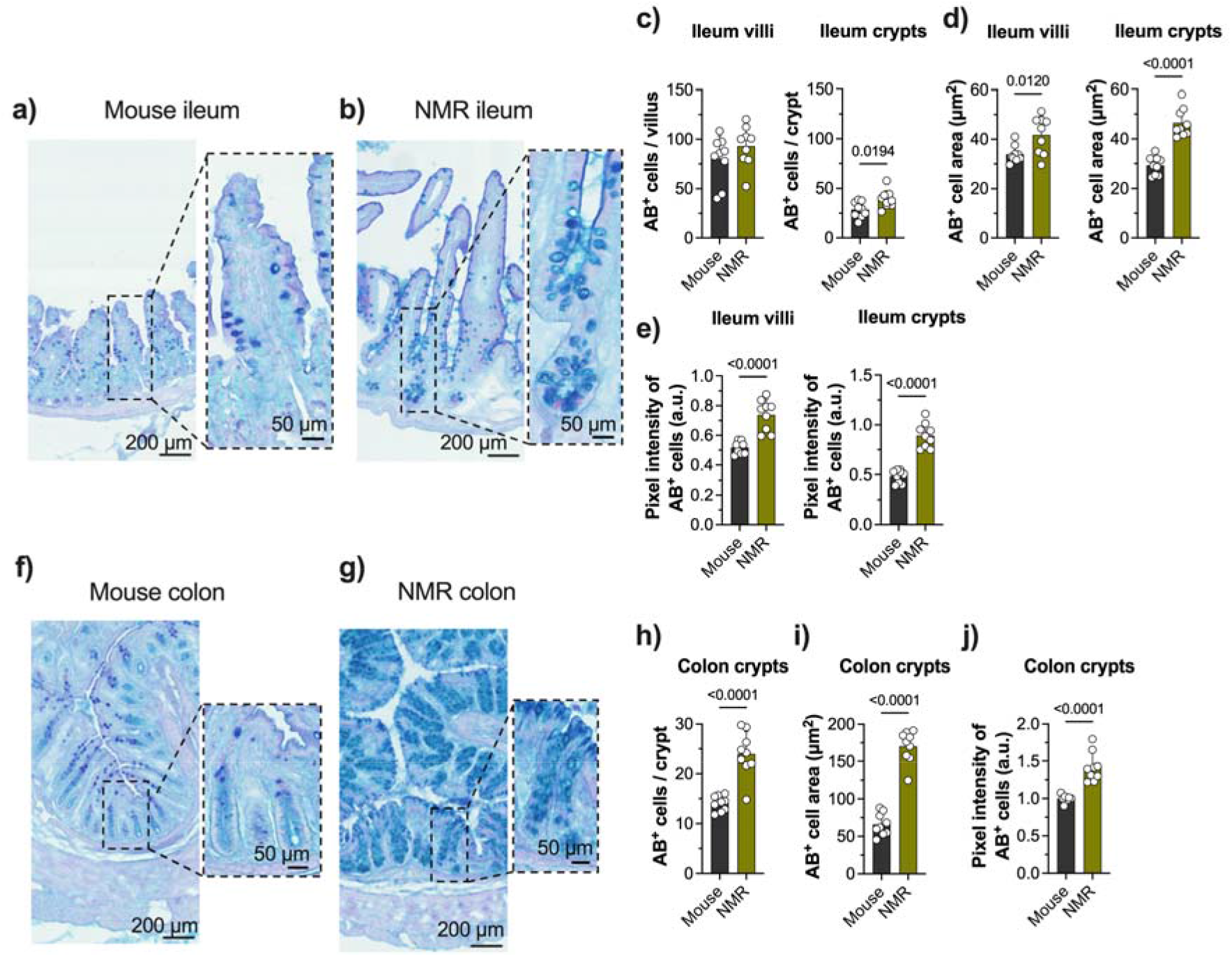
Goblet cells and mucus content in the gut of NMRs. Representative images of mouse (**a**) and NMR (**b**) ileum samples stained with AB/PAS. **c**, number of AB^+^ cells (goblet cells) in the ileum villi (left) and crypts (right) in mice (n = 9, 3 sections/animal, 3 mice) and NMRs (n = 9, 3 sections/animal, 3 NMRs). **d**, area of AB^+^ cells in the ileum villi (left) and crypts (right) in mice (n = 9, 3 sections/animal, 3 mice) and NMRs (n = 9, 3 sections/animal, 3 NMRs). **e**, pixel intensity measurement AB/PAS staining in the ileum villi (left) and crypts (right) in mice (n = 9, 3 sections/animal, 3 mice) and NMRs (n = 9, 3 sections/animal, 3 NMRs). Representative images of mouse (**f**) and NMR (**g**) colon samples stained with AB/PAS **h**, number of AB^+^ cells in the colon crypts in mice (n = 9, 3 sections/animal, 3 mice) and NMRs (n = 9, 3 sections/animal, 3 NMRs). **i**, area of AB^+^ cells in crypts in mice (n = 9, 3 sections/animal, 3 mice) and NMRs (n = 9, 3 sections/animal, 3 NMRs). **j**, pixel intensity measurement AB/PAS staining in colon crypts in mice (n = 9, 3 sections/animal, 3 mice) and NMRs (n = 9, 3 sections/animal, 3 NMRs). *P* values are shown in plots. **c**, **d**, **e**, **h**, **i** and **j**, unpaired t-test (data shown as mean ± SD).

Altogether, these findings indicate that, despite having a shorter small intestine, the NMR ileum exhibits features that maximise the surface area for nutrient and water absorption, and ionic transport, such as ∼2.5-fold larger villi compared to mice (Cohen’s d size effect = 5.3). Additionally, the larger crypts found in NMRs may be reflective of differences in tissue turnover in these animals compared to mice. Furthermore, NMR show increased numbers of larger goblet cells both in the ileum and in the colon, with a significantly higher mucus content compared to murine goblet cells. This likely confers enhanced protection upon these animals against the infiltration of microorganisms, digestive enzymes and acids, digested food particles, as well as luminal by-products and toxins.

### 3.3. Intestinal integrity: ileum and colon

Because of the clear differences in NMR intestinal structures compared to those of mice, we next aimed to examine the physical properties of this tissue. To this end, we used the gold standard technique for this purpose, i.e., the Ussing chamber for measuring mucosal membrane properties. We measured the transepithelial electrical resistance (TEER), as a measure of intestinal barrier integrity. High TEER values are indicative of decreased permeability to ions. Interestingly, the NMR terminal ileum showed a remarkably higher TEER, i.e. reduced permeability, compared to the mouse terminal ileum (mouse 45.41 ± 19.71 Ω · cm^2^ vs. NMR 145.8 ± 100.8 Ω · cm^2^, *p* = 0.0001; Cohen’s *d* size effect = 6.27; Figure 4a, b). The NMR distal colon also showed reduced permeability compared to the mouse distal colon, but the difference was less pronounced (mouse 52.35 ± 20.27 Ω · cm^2^ vs. NMR 65.65 ± 24.63 Ω · cm^2^, *p* = 0.0062; Cohen’s *d* size effect = 1.76; Figure 4c, d). While the terminal ileum in mice exhibited greater permeability compared to their colon (ileum 45.41 ± 19.71 Ω · cm^2^ vs. colon 52.35 ± 20.27 Ω · cm^2^, *p* = 0.0197; Cohen’s *d* size effect = 1.24; Supplementary Figure 2a), which aligns with prior studies in mice (*49*) and is also noted in rats (*50*, *51*) and humans (*51*, *52*), the terminal ileum of NMRs was extraordinarily less permeable than their distal colon (ileum 145.8 ± 100.8 Ω · cm^2^ vs. colon 65.65 ± 24.63 Ω · cm^2^, *p* < 0.0001; Cohen’s *d* size effect = 4.63; Supplementary Figure 2b). Altogether, these results indicate that the integrity of the gastrointestinal tract in NMRs is increased compared to mice, with the NMR small intestine (ileum) showing a strikingly low tissue permeability.

**Figure 4.**
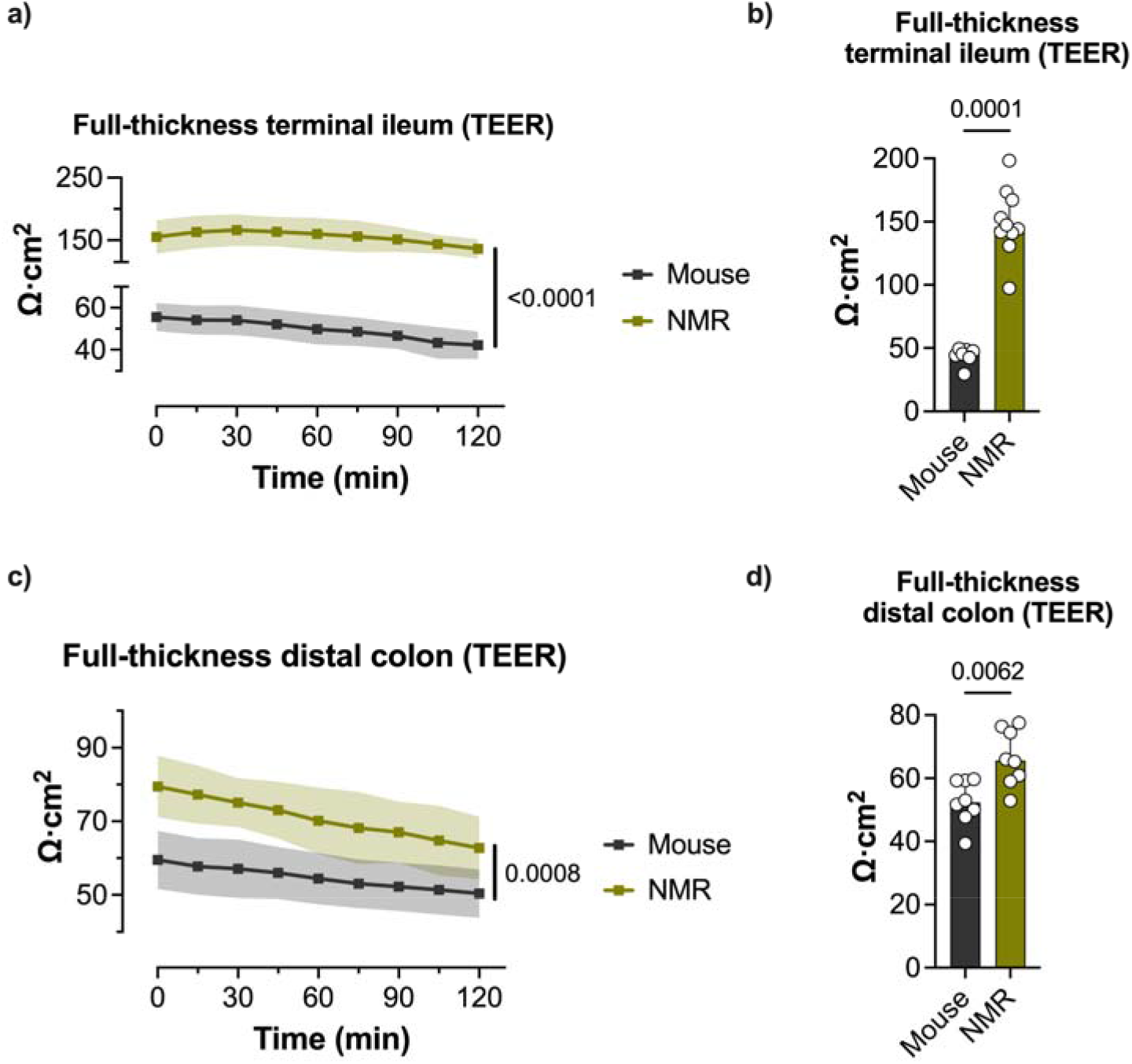
Increased intestinal integrity in NMR. **a, b)** transepithelial electrical resistance (TEER) in the terminal ileum of mice (n = 7) and NMRs (n = 10). **c**, **d)** TEER in the distal colon of mice (n = 8) and NMRs (n = 8). Data in **b** and **d** were calculated as the average TEER values between 60 min and 120 min and plotted as individual values. *P* values are shown in plots. **a**, **c**, 2-way ANOVA and Šidak’s multiple comparison test (data shown as mean, shadow area represents the SD); **b**, **d**, Mann-Whitney test (data shown as median ± IQR).

### 3.4. Ion transport in the terminal ileum

Given the different mucosal integrity properties observed in NMR vs. mouse ileal tissue, we next utilised this region to assess intestinal ion secretion. To this end, we measured the short-circuit current (*I_sc_*), as this reflects the net ionic transport, mainly Cl^-^ and Na^+^ in the small intestine, through the mucosa (*6*). An increase in *I_sc_* is primarily attributable to ion secretion to the intestinal lumen. We first selected four compounds known to evoke ion secretion in rodents and/or humans, such as 5-hydroxytryptamine (5-HT, serotonin) (*53*, *54*), bradykinin (*55*, *56*), histamine (*57*, *58*) and capsaicin (*59*, *60*). In keeping with previous studies, we confirmed that 5-HT, bradykinin, histamine, and capsaicin evoke a prominent ion secretion to the ileal mucosa in mice (Figure 5a-d). By contrast, ileum preparations from NMRs showed markedly reduced responses to 5-HT (mouse 179.8 ± 58.15 ΔμA · cm^-2^ vs. NMR 13.23 ± 17.41 ΔμA · cm^-2^, *p* < 0.0001; Figure 5a) and bradykinin (mouse 78.64 ± 22.04 ΔμA · cm^-2^ vs. NMR 17.98 ± 17.05 ΔμA · cm^-2^, *p* < 0.0001; Figure 5b), and were virtually unresponsive to histamine (mouse 22.28 ± 24.93 ΔμA · cm^-2^ vs. NMR 3.08 ± 3.16 ΔμA · cm^-2^, *p* = 0.0006; Figure 5c) and capsaicin (mouse 42.56 ± 20.20 ΔμA · cm^-2^ vs. NMR -0.74 ± 1.98 ΔμA · cm^-2^, *p* = 0.0002; Figure 5d).

**Figure 5.**
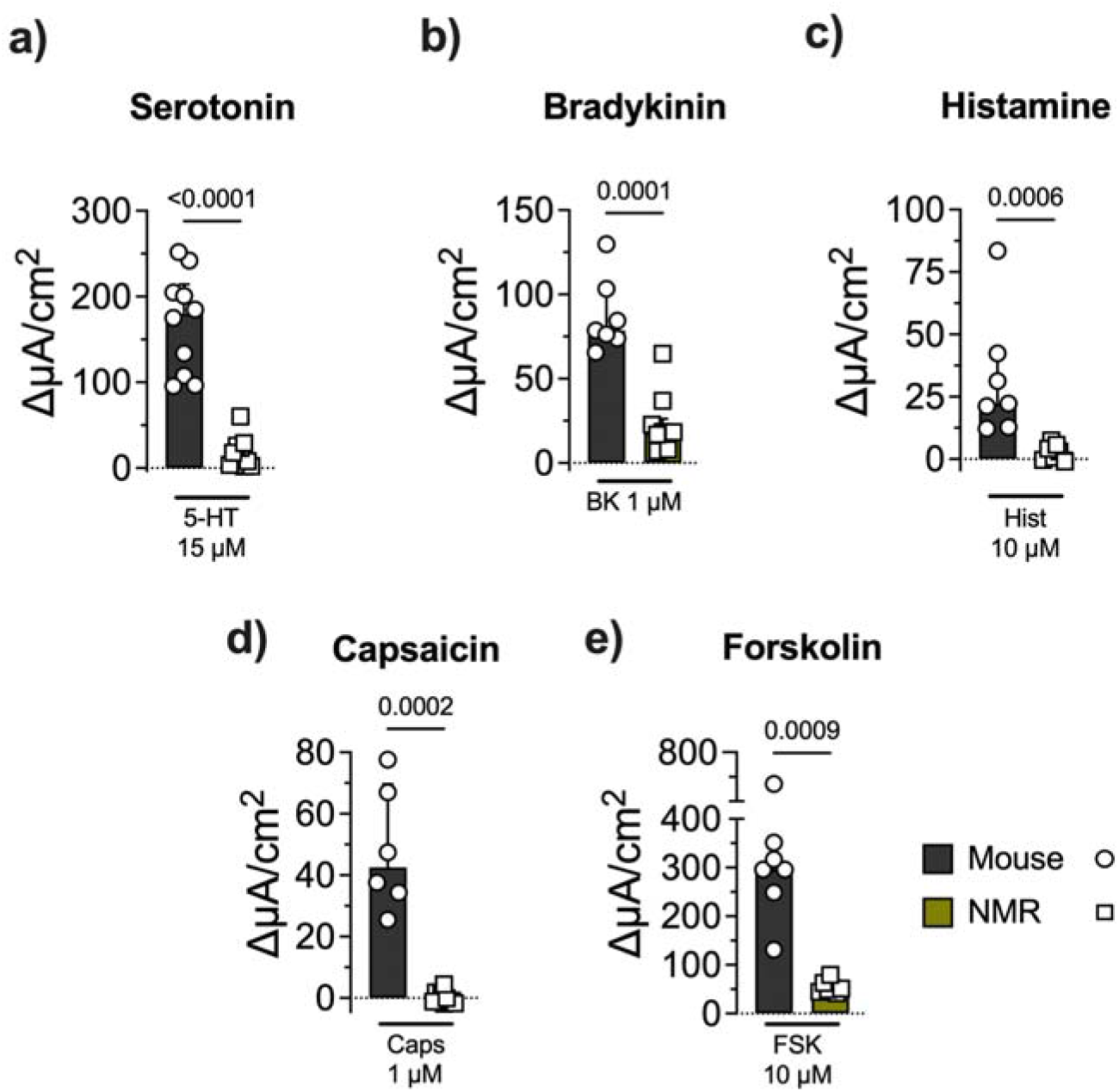
Reduced ion secretion responses in NMRs. **a)** Short-circuit current increase (Δ*Isc*) in response to 5-HT in mice (n = 10 ileum sections, N = 3 animals) and NMRs (n = 11 ileum sections, N = 3 animals). **b**) Δ*I_sc_* in response to bradykinin in mice (n = 10 ileum sections, N = 3 animals) and NMRs (n = 11 ileum sections, N = 3 animals). **c**) Δ*I_sc_* in response to histamine in mice (n = 7 ileum sections, N = 3 animals) and NMRs (n = 7 ileum sections, N = 2 animals). **d**) Δ*I_sc_* in response to capsaicin in mice (n = 6 ileum sections, N = 3 animals) and NMRs (n = 10 ileum sections, N = 3 animals). **e**) Δ*Isc* in response to forskolin in mice (n = 7 ileum sections, N = 2 animals) and NMRs (n = 7 ileum sections, N = 2 animals). For clarity, circles refer to mouse samples and squares to NMRs samples. P values are shown in plots. All data were analysed with Mann-Whitney test (data shown as median ± IQR).

Next, we aimed to further evaluate the pro-secretory capacity of the NMR ileum via receptor-independent pathways. To this end, we used the non-selective adenylate cyclase activator forskolin, which elevates the intracellular cyclic adenosine monophosphate (cAMP) concentration (*61*) and leads to increased intestinal ion secretion (Figure 5e) (*62*). Remarkably, NMR ileum tissue showed significantly reduced forskolin-evoked responses compared to mice (mouse 330.1 ± 166.1 ΔμA · cm^-2^ vs. NMR 54.41 ± 12.96 ΔμA · cm^-2^, *p* = 0.0009; Figure 5e).

Altogether, these results suggest that the cAMP signalling pathway is altered in the intestinal epithelium of NMRs, likely influencing their intestinal ion pro-secretory pathways. Moreover, NMRs display a pronounced resistance to intestinal ion secretion in response to some neurotransmitters, inflammatory mediators, and chemical irritants.

### 3.5. Expression of receptors involved on intestinal ion transport

Because NMRs showed altered ion secretion responses to 5-HT, bradykinin, histamine, and capsaicin, we next asked whether NMRs express receptors for these compounds in their intestinal tract. To this end, we used RT-qPCR to evaluate the gene expression of ion channels and receptors associated to these agonists. Importantly, some of these substances can activate several receptors (such as 5-HT, bradykinin, and histamine). Hence, we selected genes that encode receptors known to be involved in intestinal ion secretion in humans and/or rodents. Thus, we assessed the gene expression of the 5-HT receptor 4 (*Htr4*) (*53*), the bradykinin receptor B_2_ (*Bdkrb2*) (*55*), histamine H_1_ receptor (*Hrh1*) (*58*) and the capsaicin receptor, transient receptor potential vanilloid 1 (*Trpv1*). Notably, both the terminal ileum and the distal colon of NMR expressed all the genes evaluated. Ileum samples from NMR showed a reduced gene expression compared to mouse of *Bdkrb2* (mouse 0.014 ± 0.039 relative expression vs. NMR 0.001 ± 0.007 relative expression, *p* = 0.0002; Figure 6) and *Hrh1* (mouse 0.11 ± 0.73 relative expression vs. NMR 0.007 ± 0.022 relative expression, *p* = 0.0003; Figure 6), whereas similar levels of these genes were expressed in the colon of both species (*Bdkrb2*: mouse 0.01 ± 0.02 relative expression vs. NMR 0.01 ± 0.07 relative expression, *p* = 0.7984; *Hrh1*: mouse 0.01 ± 0.008 relative expression vs. NMR 0.01 ± 0.005 relative expression, *p* = 0.2220; Figure 6). By contrast, colon tissue of NMRs displayed a lower expression of *Htr4* compared to mice (mouse 0.14 ± 0.33 relative expression vs. NMR 0.01 ± 0.03 relative expression, *p* = 0.0002; Figure 6), although no such statistical difference was observed in the ileum across species for *Htr4* (mouse 0.04 ± 0.26 relative expression vs. NMR 0.18 ± 0.42 relative expression, *p* = 0.2785; Figure 6). These results indicate that, while changes in the gene expression of these receptors could partially account for the altered intestinal ion secretion observed in response to some of the compounds evaluated, NMRs express receptors for all these agents along their gastrointestinal tract. Further studies are needed to characterise intestinal secretory responses in these animals.

**Figure 6.**
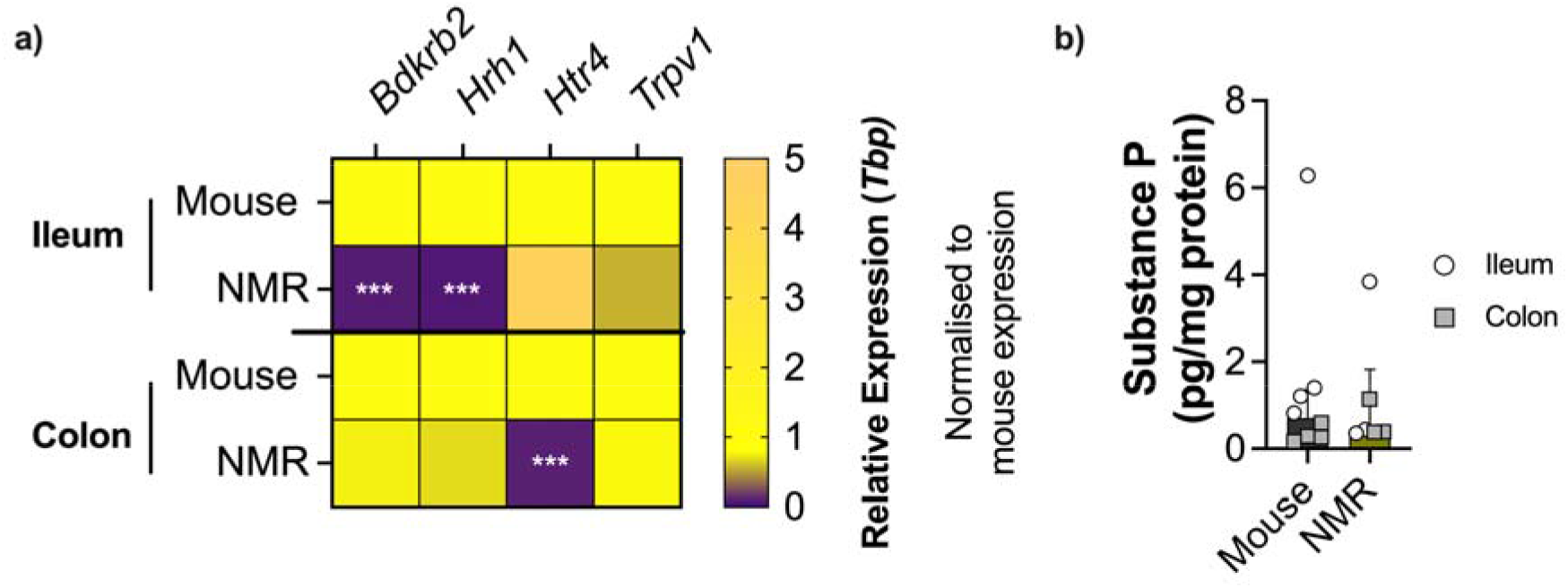
Intestinal gene expression of receptors associated with increased ion secretion and intestinal secretion of substance P. **a**) Heatmap of the relative gene expression in terminal ileum and distal colon samples in mice (n = 8) and NMRs (n = 8). **b**) Intestinal levels of substance P (normalised by the content of total protein) in mouse (n = 8; 4 ileum samples and 4 colon samples) and NMR (n = 6; 3 ileum samples and 3 colon samples). White circles refer to terminal ileum samples and grey squares to colon samples. All data were analysed with Mann-Whitney test (in **a**, data shown as colour gradient scale of the relative expression; and, in **b**, data shown as median ± IQR). Each row in **a** represents the median of the side-by-side replicates. \*\*\**P* < 0.001.

### 3.6. Substance P levels in the gut of naked mole-rats

Finally, because of its role on intestinal ion secretion (*63*, *64*), we aimed to evaluate the expression of the neuropeptide substance P in the gut of NMRs. Of interest, NMRs do not express substance P and calcitonin gene-related peptide (CGRP) in cutaneous C-fibres (*65*), in contrast to other mammalian species, including mice. Nevertheless, NMRs do produce both substance P and CGRP in longitudinal lanceolate endings associated with follicle-sinus complexes and body hair follicles (*65*), likely through Aβ-fibres. In keeping with this, we previously observed CGRP^+^ fibres in myenteric neurons in the colorectum of NMR (*43*). Therefore, we asked whether substance P is also synthetised in the NMR gut. To this end, we collected supernatants from intestinal explants (terminal ileum and distal colon) and analysed the presence of substance P by enzyme immunoassay. The levels of substance P under basal conditions were similar between NMRs and mice (mouse 1.38 ± 2.033 pg substance P · mg protein^-1^ vs. NMR 1.09 ± 1.38 pg substance P · mg protein^-1^, *p* > 0.9999; Figure 6b). Further studies will investigate the capacity of different enteric cells to produce this neuropeptide (such as neurons or enterochromaffin cells) and how this molecule and other neuropeptides modulate the gut physiology of NMRs.

## Discussion

In the present study, we investigated some aspects of the intestinal barrier function in NMRs as a potential emergent animal model in gastroenterology. Intestinal barrier dysfunction is considered one of the potential underlying pathophysiological mechanisms of many gastrointestinal diseases (*6*), and thus deemed to be of great biological and clinical relevance (*11*). The persisting incidence and prevalence of gastrointestinal conditions in the last decades (*1*) reflects the unmet need for novel strategies in the field. Here, we used NMRs as a non-traditional animal model due to their exceptional biological features (*66*) and studied their intestinal physiology. We found that the NMR small intestine was significantly shorter compared to that of mice, although it displayed a more robust tissue architecture (thicker intestinal layers, longer ileum villi, bigger crypts, and elevated mucus content). In keeping with this, the integrity (measured as TEER) of the NMR ileum was remarkably increased compared to the mouse ileum. The colon of NMR, on the other hand, showed more structural similarities to mice, but again displayed enhanced mucus content and superior tissue integrity. Of particular interest, we found that the ileum of NMRs showed reduced ion secretion responses to 5-HT, bradykinin, histamine, and capsaicin, even though receptors for these agonists were expressed in the gastrointestinal tract. Taken together, NMRs represent an animal model of considerable interest for gastroenterology and, particularly, for studying intestinal barrier function.

We first characterised the morphoanatomical features of the intestinal tract in NMRs. The NMR’s notably shorter small intestinal length has likely led to the simultaneous development of longer villi compared to those mice, or rats (*67*). NMRs do not drink water and fulfil their hydration requirements entirely through their solid diet. Therefore, we presume that longer villi enable NMRs to increase the surface area to volume ratio for nutrient and water absorption, which has led to water consumption being redundant. In keeping with this, a recent study showed that long villi are found along the entire small bowel of NMR (*23*), as well as enterocytes involved in the digestion and absorption of nutrients and water (*23*, *68*). Furthermore, NMRs exhibited a greater number of goblet cells in the ileum and colon. Additionally, the size of these cells was substantially larger in NMRs and had increased mucus granule content compared to mice. These results strongly suggest that goblet cells in NMRs have an enhanced capacity to produce mucus/mucins. In keeping with this, it was recently reported that NMRs have a thicker mucus layer and elevated expression of *Muc2* (gene encoding for the oligomeric mucus gel-forming protein Mucin 2) compared to mice (*23*), likely conferring an increased protection against environmental chemicals (*23*).

Notably, the histological features of the NMRs small intestine and colon more closely resemble those of bigger rodents, such as rats (*69*) and guinea pigs (*70, 71*). This is of particular interest because the altered intestinal architecture found in NMRs would not completely explain the increased ileum tissue integrity (reflected as increased in TEER) observed in these animals compared to mice. In fact, previous studies utilising Ussing chambers have reported >2-fold lower TEER values in the ileum of guinea pigs (*72*) and rats (*73, 74*) compared to our results in NMRs, and similar values in the colon of rats (*73–75*) compared to our findings. This strongly suggests that the small intestine of NMRs possesses strong tight junction structures that increase the integrity of this barrier. It is important to note, however, that flux across tight junctions occurs by two distinct routes, referred to as the “pore” and “leak” pathways. In the current study, we measured TEER as a proxy for intestinal integrity, but this does not discriminate between these two different pathways (*6*). The pore pathway is a high-capacity route that selectively allows passage of molecules based on charge and size, with a maximum diameter of ∼4-6 Å. On the other hand, the low-capacity leak pathway is less selective and allows the passage of larger molecules (up to ∼100 Å), irrespective of the charge. Future studies are needed to evaluate in detail differences in the pore and leak pathways between the small intestine and colon of NMRs. Moreover, due to the crucial role of different tight junctions to selectively regulate each of these pathways, the regulation of these barrier components is of great relevance. Thus, future studies will assess the expression, organisation and interprotein interactions of tight junctions in the NMR small intestine and how these control the pore and leak pathways. Overall, our data support that the specific characteristics of the NMR small intestine may represent an interesting model for studying intestinal barrier dysfunction in coeliac disease (*76*) and irritable bowel syndrome (*77*), amongst other conditions(*23*).

Another crucial function of the intestinal barrier is the transport of ions and fluids across the epithelium. Disruption of this well-coordinated process may lead to diarrhoea when the gut’s capacity for fluid absorption is exceeded. Of note, we found that the NMR ileum displays reduced ion secretion responses to four different compounds: 5-HT, bradykinin, histamine, and capsaicin. The bradykinin result aligns with our prior observation that this inflammatory mediator fails to activate colonic afferent nerves in NMRs (*43*). However, we did observe that bradykinin led to mechanical sensitisation of these fibres (*43*), suggesting that they express functional bradykinin B_2_ receptors. In the present study, we confirmed that NMRs indeed express *Bdkrb2* (gene encoding the bradykinin B_2_ receptor) in both the colon and ileum. However, differences in the functionality of these receptors and the downstream effector signalling in NMRs remain to be elucidated. Future studies will also explore the different cells expressing this marker (epithelial cells (*78*) and nerve fibres (*79*)) in NMRs, and how these modulate visceral pain and mucosal ion secretion. The role of 5-HT in the NMR gastrointestinal tract has not been previously characterised. While fibroblasts isolated from the back skin or the lungs of NMRs express high levels of 5-HT (*80*), plasma circulating levels of this neurotransmitter are significantly lower compared to mice (*81*). In the gut, we showed that the ion secretion induced by 5-HT in the ileum was decreased in NMRs compared to mice, although the expression of *Htr4* in this tissue was similar in both species. Further studies are therefore required to determine the functionality of this and other serotonergic receptors, such as 5-HT receptors 1, 2 and 3 (*82*), in different cell types in the NMR intestine. Finally, the lack of cutaneous neuropeptides and unusual connectivity of cutaneous afferent neurons in the spinal cord in NMRs have been proposed to explain the absence capsaicin-induced nocifensive behaviour and associated thermal hypersensitivity (*83*), as well as the lack of histamine-induced scratching behaviour (*84*) in these animals. However, it should be noted that the lack of substance P and CGRP alone cannot completely explain the lack of pain- and itch-associated behaviours induced by these compounds, as *Tac1*-*Calca* double knockout mice (therefore lacking both substance P and CGRP peptides) display intact responses to mechanical, thermal, chemical, visceral pain, itch, and neurogenic inflammation (*85*). In the present study, we showed that despite the virtually complete absence of ion secretion induced by histamine and capsaicin in NMRs, these animals display intestinal basal levels of substance P comparable to mice. Future studies will investigate the role of substance P and other neurotransmitters/neuropeptides in the gut physiology of NMRs, as well as their cellular sources. Overall, we propose that NMRs may be a promising model for the pathogenesis of inflammation-associated diarrhoea in inflammatory bowel disease (*86*) and serotonin- and histamine-associated diarrhoea in irritable bowel syndrome (*87, 88*).

The most notable observation regarding intestinal ion secretion is the reduced response to the cAMP pathway-activator forskolin observed in NMR tissue. cAMP is a ubiquitous second messenger found across all living domains that is generated by adenylate cyclases upon conversion of adenosine triphosphate (ATP) to cAMP (*89*). Forskolin activates all adenylate cyclases except AC9 and sAC (*90*), dramatically increasing the levels of cAMP in virtually all tissues (*61*). Our findings raise numerous questions, not only concerning the intestinal ion pro-secretory pathways of NMRs, but also more broadly about their cell biology. In the context of intestinal ion secretion, the conventional paradigm indicates that activation of the cAMP/protein kinase A (PKA) pathway activates the cystic fibrosis transmembrane conductance regulator (CFTR) protein, representing the primary route for epithelial ion secretion (*91, 92*). Of interest, a study (*93*) showed that adenylate cyclase 6 (AC6) is the most abundant isoform in the gut of mouse and humans and is physically and functionally associated with CFTR at the apical surface of intestinal epithelial cells. Interestingly, these authors found that gut epithelial cells lacking AC6 fail to generate cAMP in response to forskolin and AC6-deficient tracheal cells showed reduced forskolin-evoked Cl^−^ secretion compared to preparations from littermate controls (*93*). Moreover, AC6-deficient mice were protected from cholera toxin-induced diarrhoea (*93*), suggesting an important role of this isoform in ion and water secretion. Whether AC6 is present in the NMR gut or if the functionality of this and other adenylate cyclases is altered in this species remains to be investigated. Furthermore, the functionality of the CFTR protein and associated signalling pathways in this species should be investigated in detail. Finally, given that the cAMP system plays fundamental roles in the regulation of the signalling pathways of serotonin (*94*), bradykinin (*95*), and TRPV1 (*96*), this pathway could also interfere with the mechanisms leading to receptor-dependent ion secretion studied here. Therefore, further studies are needed to characterise the cAMP signalling system in NMRs.

We recognise that our study has some limitations. Although approximately physiologically age-matched animals for both species were used, intestinal permeability typically changes throughout the lifetime. However, it should be noted that differences in intestinal permeability between mice and NMRs are in the opposite direction of what one would expect, that is, an increased permeability in older animals (*97*). Interestingly, we found that the ileum permeability was similar in young NMRs compared to older animals (young 153.0 ± 57.09 Ω · cm^2^ vs. older 144.2 ± 42.75 Ω · cm^2^, *p* = 0.7000; Supplementary Figure 2c), thus demonstrating that the differences observed between species are not related to age. Furthermore, all NMRs used in the present study were non-breeders and thus we cannot comment on whether or findings would hold true in sexual mature, breeding animals. Sexual maturity is of relevance to our work because it has been shown, for example, that progesterone decreases gut permeability in humans and Caco-2 cells (*98*) and that oestrogen modulates ion secretion in intestinal epithelial cells (*99*). Most of the NMRs used in our study were males (only 1 female was used in TEER experiments and another 1 in short-circuit current experiments), and thus we are unable to draw firm conclusions with regard to any potential sex differences, but being non-breeder animals, all were sexually immature. However, we do not believe that sex and/or state of sexual maturity are likely responsible for the significant differences observed in the NMR ileum, as the relevant hormones would similarly affect the colon, where we found similar TEER values as previously reported in rats (*73–75*). It should be noted as well that the Ussing chambers experiments were performed at 37°C, that is, 5-7°C above the NMRs physiological temperature (*100*). However, an increase in the physiological temperature is typically associated with an increase in mucosal permeability (*101, 102*). It remains to be elucidated whether changes in physiological temperature affect intestinal function in NMRs, although one would normally expect changes in the opposite direction to our findings. Finally, other caveats that must be considered are the differences in diet and microbiota between mice and NMRs, which can affect both the intestinal integrity (*103*) and the ion secretion (*104*). Whether specific gut bacteria found in NMRs (*25*) or compounds of their diet are important to improve intestinal barrier function remain to be elucidated.

In conclusion, our work illustrates a remarkably low permeability in the NMR small intestine. Moreover, it reveals that these animals have reduced intestinal ion secretion responses to 5-HT, bradykinin, histamine, and capsaicin. Overall, this study adds further evidence to the uncommon biological features exhibited by this species. The NMR gut may therefore offer particular insights into the physiology relevant to gastrointestinal conditions and aid in devising strategies for their treatments. Thus, we strongly support the use of this unconventional animal model in further investigations of intestinal barrier function.

## 4. Acknowledgements

JA-L and EStJS acknowledge funding from the Medical Research Council (MRC) (MR/W002426/1) and AR was supported by AstraZeneca PhD Studentship (G115018).

## 6. Author contribution

JA-L conceptualised and designed the research study, conducted the experiments (Ussing chamber experiments, gene expression, enzyme immunoassays), acquired and analysed the data and wrote the manuscript. AR conducted the experiments (histology, AB/PAS staining and IHC) and acquired the data. DCB provided experimental support and discussed the data. EStJS provided experimental support, reviewed, and discussed the data and revised the manuscript. All authors corrected and approved the final version of the manuscript.

## 7. Data availability

Data sets supporting the conclusions of this article are available in University of Cambridge Apollo Repository (http://doi.org/10.17863/CAM.106848).

## 10. Supplementary information

**Supplementary Table 1.**
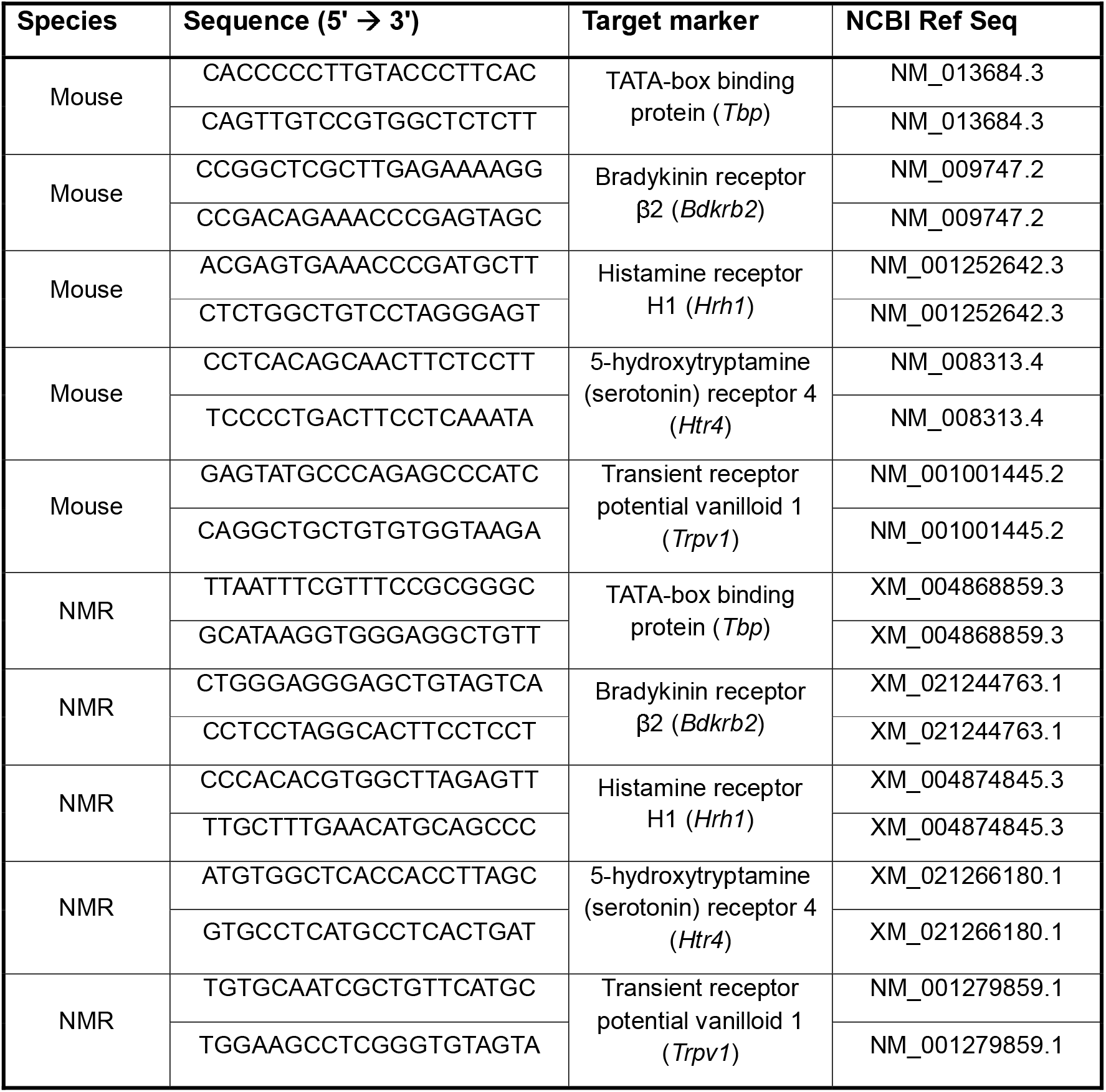
Primer sequences for gene detection by RT-qPCR.

**Supplementary Figure 1.**
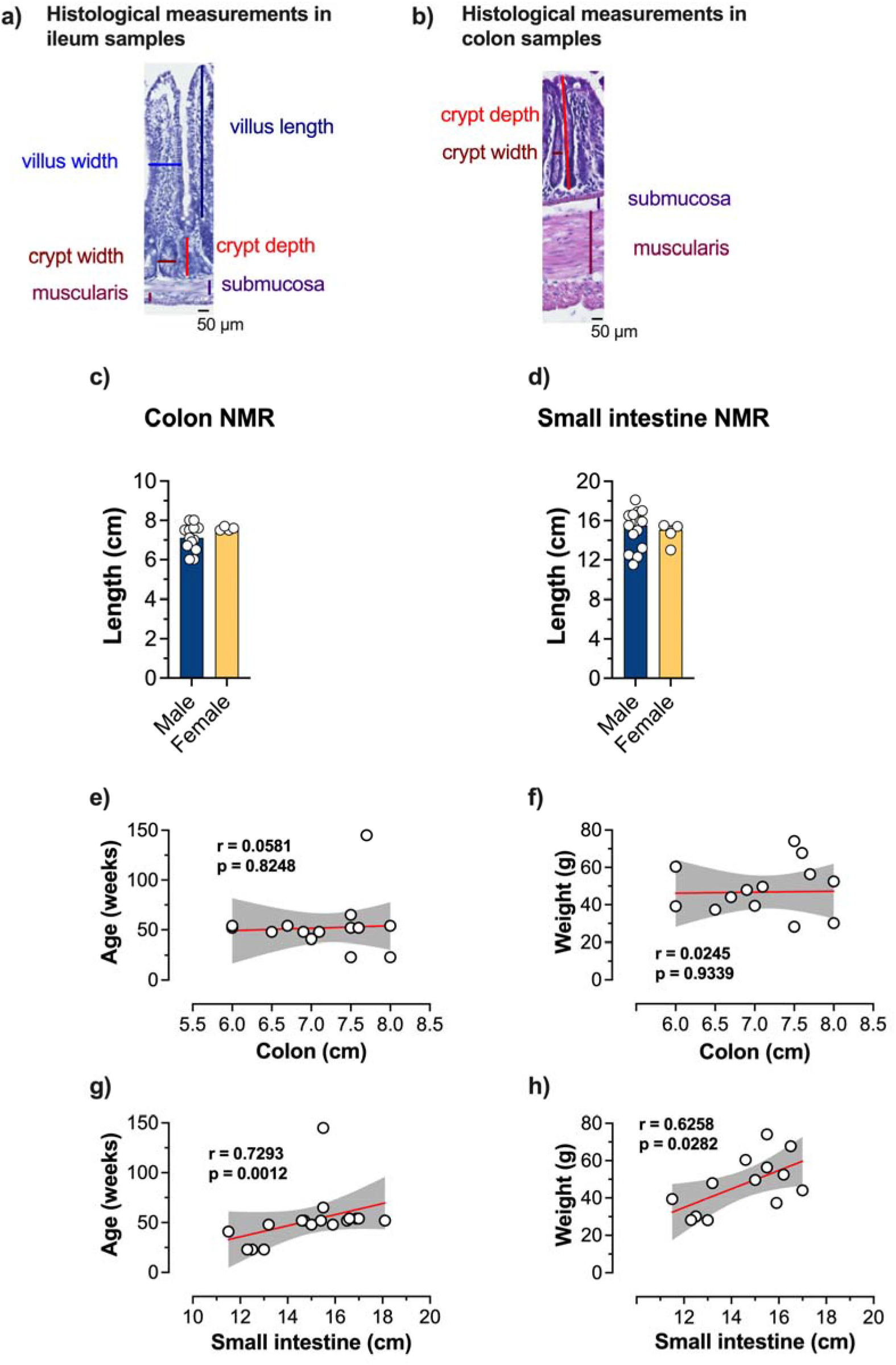
Extended anatomical analyses of the NMR gastrointestinal tract. Representative images of the histological measurements performed in ileum (**a**) and colon (**b**) samples. **c)** Colon length in male (n = 13) and female (n = 4) NMRs. **d)** Small intestine length in male (n = 13) and female (n = 4) NMRs. **e)** Correlation between age and colon length in NMRs (n = 17). **f)** Correlation between weight and colon length in NMRs (n = 17). **g)** Correlation between age and small intestine length in NMRs (n = 17). **h)** Correlation between weight and small intestine length in NMRs (n = 16). *P* values are shown in plots. **c** and **d**, Mann Whitney test (data shown as median ± IQR); **e**, **f**, **g** and **h**, two-tailed Pearson’s correlation (data shown as individual data, and 95% CI).

**Supplementary Figure 2.**
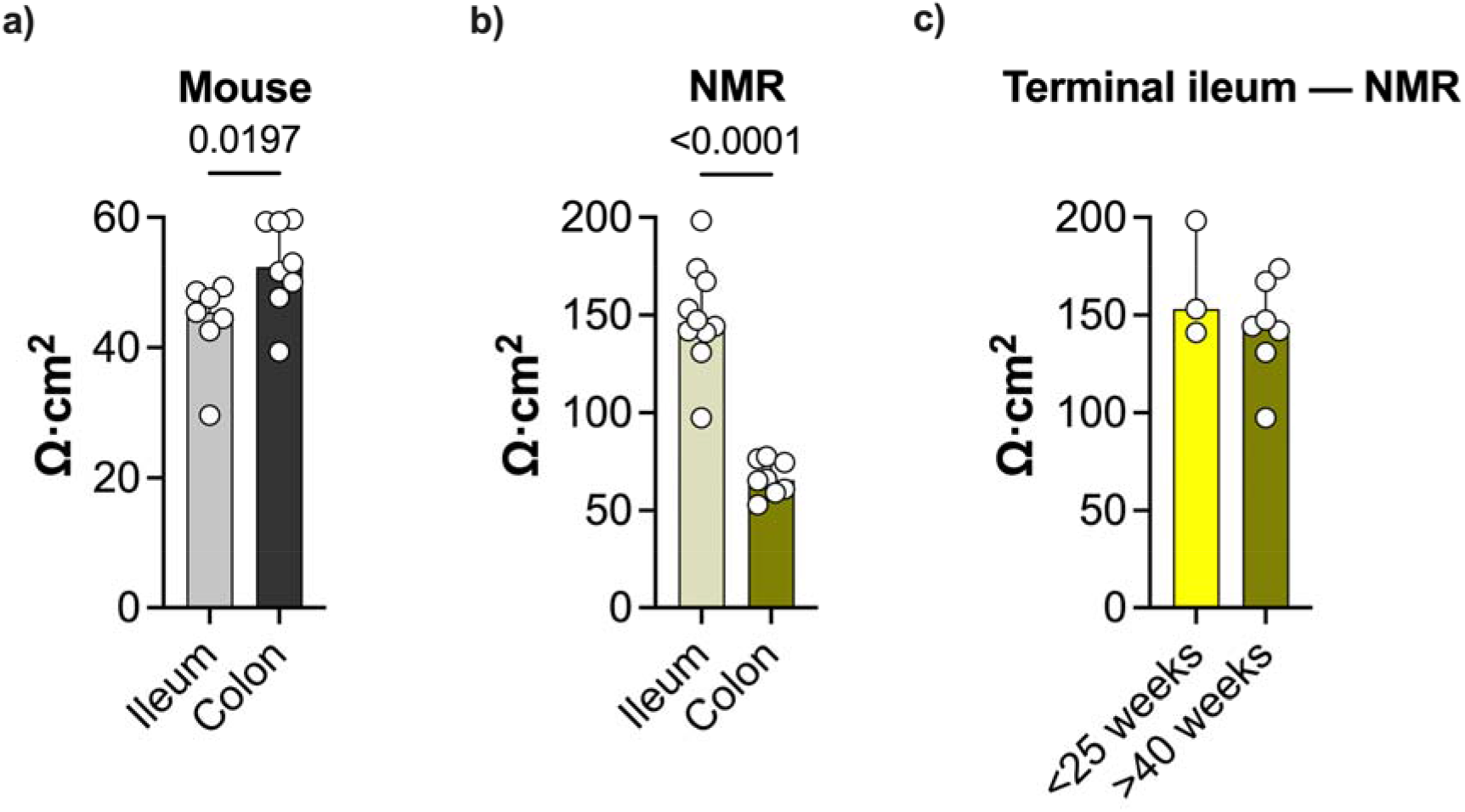
Extended analyses of the tissue integrity in NMR and mice. **a)** TEER in the terminal ileum (n = 7) and colon (n = 8) in mice. **b)** TEER in the terminal ileum (n = 10) and colon (n = 8) in NMRs. **c)** TEER in the terminal ileum of young (<25 weeks old; n = 3) and old (>40 weeks old; n = 7) NMRs. Data were calculated as the average TEER values between 60 min and 120 min and plotted as individual values. *P* values are shown in plots. **a**, **b** and **c**, Mann-Whitney test (data shown as median ± IQR).

## 9. References

1. Y. Wang, Y. Huang, R. C. Chase, T. Li, D. Ramai, S. Li, X. Huang, S. O. Antwi, A. P. Keaveny, M. Pang, Global Burden of Digestive Diseases: A Systematic Analysis of the Global Burden of Diseases Study, 1990 to 2019. Gastroenterology 165, 773–783.e15 (2023).

2. T. C. Rose, A. Pennington, C. Kypridemos, T. Chen, M. Subhani, J. Hanefeld, L. Ricciardiello, B. Barr, Analysis of the burden and economic impact of digestive diseases and investigation of research gaps and priorities in the field of digestive health in the European Region—White Book 2: Executive summary. United European Gastroenterol J 10, 657–662 (2022).

3. V. C. Goodoory, C. E. Ng, C. J. Black, A. C. Ford, Direct healthcare costs of Rome IV or Rome III-defined irritable bowel syndrome in the United Kingdom. Aliment Pharmacol Ther 56, 110–120 (2022).

4. R. S. Sandler, J. E. Everhart, M. Donowitz, E. Adams, K. Cronin, C. Goodman, E. Gemmen, S. Shah, A. Avdic, R. Rubin, The burden of selected digestive diseases in the United States. Gastroenterology 122, 1500–1511 (2002).

5. L. A. Pattison, A. Cloake, S. Chakrabarti, H. Hilton, R. H. Rickman, J. P. Higham, M. Y. Meng, L. W. Paine, L. Qiu, A. Ritoux, D. C. Bulmer, G. Callejo, E. St John Smith, Digging deeper into pain – an ethological behavior assay correlating well-being in mice with human pain experience. Pain, doi: 10.1101/2023.08.18.553862 (2024).

6. T. Vanuytsel, J. Tack, R. Farre, The Role of Intestinal Permeability in Gastrointestinal Disorders and Current Methods of Evaluation. Front Nutr 8, 717925 (2021).

7. H. Majamaa, E. Isolauri, Evaluation of the gut mucosal barrier: Evidence for increased antigen transfer in children with atopic eczema. J Allergy Clin Immunol 97, :985–90 (1996).

8. A. Benard, P. Desreumeaux, D. Huglo, A. Hoorelbeke, A.-B. Tonnel, B. Wallaert, iD Lille, Increased intestinal permeability in bronchial asthma. J Allergy Clin Immunol 97, 1173–1181 (1996).

9. A. Fasano, N. P. Visanji, L. W. C. Liu, A. E. Lang, R. F. Pfeiffer, Gastrointestinal dysfunction in Parkinson’s disease. Lancet Publishing Group [Preprint] (2015). 10.1016/S1474-4422(15)00007-1.

10. M. P. Fink, Leaky gut hypothesis: a historical perspective. [Preprint] (1990). 10.1097/00003246-199005000-00024.

11. M. Camilleri, Leaky gut: mechanisms, measurement and clinical implications in humans. BMJ Publishing Group [Preprint] (2019). 10.1136/gutjnl-2019-318427.

12. H. Vanheel, M. Vicario, T. Vanuytsel, L. Van Oudenhove, C. Martinez, À. V. Keita, N. Pardon, J. Santos, J. D. Soderholm, J. Tack, R. Farré, Impaired duodenal mucosal integrity and low-grade inflammation in functional dyspepsia. Gut 63, 262–71 (2014).

13. A. Fedi, C. Vitale, G. Ponschin, S. Ayehunie, M. Fato, S. Scaglione, In vitro models replicating the human intestinal epithelium for absorption and metabolism studies: A systematic review. Journal of Controlled Release 335, 247–268 (2021).

14. J. Fogh, T. Orfeo, J. Tiso, F. E. Sharkey, Establishment of human colon carcinoma lines in nude mice. Exp Cell Biol 47, 136–144 (1979).

15. M. González-González, C. Díaz-Zepeda, J. Eyzaguirre-Velásquez, C. González-Arancibia, J. A. Bravo, M. Julio-Pieper, Investigating gut permeability in animal models of disease. Frontiers Media S.A. [Preprint] (2019). 10.3389/fphys.2018.01962.

16. J. Foulke-Abel, J. In, J. Yin, N. C. Zachos, O. Kovbasnjuk, M. K. Estes, H. De Jonge, M. Donowitz, Human Enteroids as a Model of Upper Small Intestinal Ion Transport Physiology and Pathophysiology. Gastroenterology 150, 638–649.e8 (2016).

17. F. Weiß, D. Holthaus, M. Kraft, C. Klotz, M. Schneemann, J. D. Schulzke, S. M. Krug, Human duodenal organoid-derived monolayers serve as a suitable barrier model for duodenal tissue. Ann N Y Acad Sci 1515, 155–167 (2022).

18. A. J. Barker, U. Koch, G. R. Lewin, S. J. Pyott, The Extraordinary Biology of the Naked Mole-Rat (2021)vol. 1319.

19. R. Buffenstein, The Naked Mole-Rat: A New Long-Living Model for Human Aging Research. J Gerontol A Biol Sci Med Sci 60, 1369–77 (2005).

20. K. Oka, M. Yamakawa, Y. Kawamura, N. Kutsukake, K. Miura, The Naked Mole-Rat as a Model for Healthy Aging. Annu Rev Anim Biosci 11, 207–226 (2023).

21. G. Maldonado, G. Hernández, Translational control in the naked mole-rat as a model highly resistant to cancer. [Preprint] (2021). 10.1016/j.bbcan.2020.188455.

22. Y. H. Edrey, D. X. Medina, M. Gaczynska, P. A. Osmulski, S. Oddo, A. Caccamo, R. Buffenstein, Amyloid beta and the longest-lived rodent: the naked mole-rat as a model for natural protection from Alzheimer’s disease. Neurobiol Aging 34, 2352–60 (2013).

23. S. Montazid, S. Bandyopadhyay, D. W. Hart, N. Gao, B. Johnson, S. G. Thrumurthy, D. J. Penn, B. Wernisch, M. Bansal, P. M. Altrock, F. Rost, P. Gazinska, P. Ziolkowski, B. H. Hayee, Y. Liu, J. Han, A. Tessitore, J. Koth, W. F. Bodmer, J. E. East, N. C. Bennett, I. Tomlinson, S. Irshad, Adult stem cell activity in naked mole rats for long-term tissue maintenance. Nat Commun 14, 8484 (2023).

24. S. Wirtz, V. Popp, M. Kindermann, K. Gerlach, B. Weigmann, S. Fichtner-Feigl, M. F. Neurath, Chemically induced mouse models of acute and chronic intestinal inflammation. Nat Protoc 12, 1295–1309 (2017).

25. T. Debebe, E. Biagi, M. Soverini, S. Holtze, T. B. Hildebrandt, C. Birkemeyer, D. Wyohannis, A. Lemma, P. Brigidi, V. Savkovic, B. König, M. Candela, G. Birkenmeier, Unraveling the gut microbiome of the long-lived naked mole-rat. Sci Rep 7, 9590 (2017).

26. H. G. Hilton, N. D. Rubinstein, P. Janki, A. T. Ireland, N. Bernstein, N. L. Fong, K. M. Wright, M. Smith, D. Finkle, B. Martin-McNulty, M. Roy, D. M. Imai, V. Jojic, R. Buffenstein, Single-cell transcriptomics of the naked molerat reveals unexpected features of mammalian immunity. PLoS Biol 17, e3000528 (2019).

27. T. S. Taylor, P. Konda, S. S. John, D. C. Bulmer, J. R. F. Hockley, E. S. J. Smith, Galanin suppresses visceral afferent responses to noxious mechanical and inflammatory stimuli. Physiol Rep 8, e14326 (2020).

28. S. U. Mondelaers, S. A. Theofanous, M. V. Florens, E. Perna, J. Aguilera-Lizarraga, G. E. Boeckxstaens, M. M. Wouters, W. M. Mondelaers SU, Theofanous SA, Florens MV, Perna E, Aguilera-Lizarraga J, Boeckxstaens GE, S. U. Mondelaers, S. A. Theofanous, M. V. Florens, E. Perna, J. Aguilera-Lizarraga, G. E. Boeckxstaens, M. M. Wouters, Effect of genetic background and postinfectious stress on visceral sensitivity in Citrobacter rodentium-infected mice. Neurogastroenterology and Motility 28, 647–658 (2016).

29. P. Bankhead, M. B. Loughrey, J. A. Fernández, Y. Dombrowski, D. G. McArt, P. D. Dunne, S. McQuaid, R. T. Gray, L. J. Murray, H. G. Coleman, J. A. James, M. Salto-Tellez, P. W. Hamilton, QuPath: Open source software for digital pathology image analysis. Sci Rep 7, 16878 (2017).

30. K. R. Rogers-Broadway, E. Karteris, Amplification efficiency and thermal stability of qPCR instrumentation: Current landscape and future perspectives. Exp Ther Med 10, 1261–1264 (2015).

31. D. Svec, A. Tichopad, V. Novosadova, M. W. Pfaffl, M. Kubista, How good is a PCR efficiency estimate: Recommendations for precise and robust qPCR efficiency assessments. Biomol Detect Quantif 3, 9–16 (2015).

32. N. Eissa, H. Hussein, H. Wang, M. F. Rabbi, C. N. Bernstein, J. E. Ghia, Stability of reference genes for messenger RNA quantification by real-time PCR in mouse dextran sodium sulfate experimental colitis. PLoS One 11, e0156289 (2016).

33. N. Eissa, L. Kermarrec, H. Hussein, C. N. Bernstein, J. E. Ghia, Appropriateness of reference genes for normalizing messenger RNA in mouse 2,4-dinitrobenzene sulfonic acid (DNBS)-induced colitis using quantitative real time PCR. Sci Rep 7, 42427 (2017).

34. F. Wang, J. Wang, D. Liu, Y. Su, Normalizing genes for real-time polymerase chain reaction in epithelial and nonepithelial cells of mouse small intestine. Anal Biochem 399, 211–217 (2010).

35. J. Zhang, Y. Y. Gao, Y. Q. Huang, Q. Fan, X. T. Lu, C. K. Wang, Selection of housekeeping genes for quantitative gene expression analysis in yellow-feathered broilers. Ital J Anim Sci 17, 540–546 (2018).

36. A. B. Nygard, C. B. Jørgensen, S. Cirera, M. Fredholm, Selection of reference genes for gene expression studies in pig tissues using SYBR green qPCR. BMC Mol Biol 8 (2007).

37. K. J. Livak, T. D. Schmittgen, Analysis of relative gene expression data using real-time quantitative PCR and the 2(-Delta Delta C(T)) Method. Methods, doi: 10.1006/meth.2001.1262 (2001).

38. J. Aguilera-Lizarraga, M. V. Florens, M. F. Viola, P. Jain, L. Decraecker, I. Appeltans, M. Cuende-Estevez, N. Fabre, K. Van Beek, E. Perna, D. Balemans, N. Stakenborg, S. Theofanous, G. Bosmans, S. U. Mondelaers, G. Matteoli, S. Ibiza Martínez, C. Lopez-Lopez, J. Jaramillo-Polanco, K. Talavera, Y. A. Alpizar, T. B. Feyerabend, H. R. Rodewald, R. Farre, F. A. Redegeld, J. Si, J. Raes, C. Breynaert, R. Schrijvers, C. Bosteels, B. N. Lambrecht, S. D. Boyd, R. A. Hoh, D. Cabooter, M. Nelis, P. Augustijns, S. Hendrix, J. Strid, R. Bisschops, D. E. Reed, S. J. Vanner, A. Denadai-Souza, M. M. Wouters, G. E. Boeckxstaens, Local immune response to food antigens drives meal-induced abdominal pain. Nature 590, 151–156 (2021).

39. S. De Schepper, S. Verheijden, J. Aguilera-Lizarraga, M. F. Viola, W. Boesmans, N. Stakenborg, I. Voytyuk, I. Smidt, B. Boeckx, I. Dierckx de Casterlé, V. Baekelandt, E. Dominguez Gonzalez, M. Mack, I. Depoortere, B. De Strooper, B. Sprangers, U. Himmelreich, S. Soenen, M. Guilliams, P. Vanden Berghe, E. Jones, D. Lambrechts, G. Boeckxstaens, Self-Maintaining Gut Macrophages Are Essential for Intestinal Homeostasis. Cell 175, 400–415.e13 (2018).

40. C. Wang, R. Weindruch, J. R. Fernández, C. S. Coffey, P. Patel, D. B. Allison, Caloric restriction and body weight independently affect longevity in Wistar rats. International journal of obesity and related metabolic disorders: journal of the International Association for the Study of Obesity. 28, 357–362 (2004).

41. M. Altun, E. Bergman, E. Edström, H. Johnson, B. Ulfhake, Behavioral impairments of the aging rat. Physiol Behav 92, 911–923 (2007).

42. B. Wiggs, A. Churg, J. L. Wright, Influence of weight on pulmonary function in the adult guinea pig. Respiration 58, 37–41 (1991).

43. J. R. F. Hockley, K. H. Barker, T. S. Taylor, G. Callejo, Z. M. Husson, D. C. Bulmer, E. S. J. Smith, Acid and inflammatory sensitisation of naked mole-rat colonic afferent nerves. Mol Pain 16, 1744806920903150 (2020).

44. F. Treves, Anatomy of the Intestinal Canal and Peritoneum in Man. British Medical Journal 1, 580–583 (1885).

45. B. M. L. Underhill, Intestinal length in man. British Medical Journal 2, 1243–1246 (1955).

46. G. Hounnou, C. Destrieux, J. Desmé, P. Bertrand, S. Velut, Anatomical study of the length of the human intestine. Surgical and Radiologic Anatomy 24, 290–294 (2002).

47. S. Sadahiro, T. Ohmura, Y. Yamada, T. Saito, Y. Taki, Analysis of length and surface area of each segment of the large intestine according to age, sex and physique. Surg Radiol Anat 14, 251–257 (1992).

48. L. T. Weaver, S. Austin, T. J. Cole, M. Dunn, Small intestinal length: a factor essential for gut adaptation. Gut 32, 1321–1323 (1991).

49. N. Hanning, R. Verboven, J. G. De Man, H. Ceuleers, H. U. De Schepper, A. Smet, B. Y. De Winter, Single-day and multi-day exposure to orogastric gavages does not affect intestinal barrier function in mice. American Journal of Physiology-Gastrointestinal and Liver Physiology 324, G281–G294 (2023).

50. L. Feighery, A. Smyth, S. Keely, A. W. Baird, W. T. O’Connor, J. J. Callanan, D. J. Brayden, Increased intestinal permeability in rats subjected to traumatic frontal lobe percussion brain injury. Journal of Trauma - Injury, Infection and Critical Care 64, 131– 137 (2008).

51. P. Nejdfors, M. Ekelund, B. Jeppsson, B. R. Weström, Mucosal in vitro permeability in the intestinal tract of the pig, the rat, and man: Species- and region-related differences. Scand J Gastroenterol 35, 501–507 (2000).

52. Å. Sjöberg, M. Lutz, C. Tannergren, C. Wingolf, A. Borde, A. L. Ungell, Comprehensive study on regional human intestinal permeability and prediction of fraction absorbed of drugs using the Ussing chamber technique. European Journal of Pharmaceutical Sciences 48, 166–180 (2013).

53. M. R. Budhoo, R. P. Harris, J. M. Kellum, 5-Hydroxytryptamine-induced Cl-transport is mediated by 5-HT3 and 5-HT4 receptors in the rat distal colon. Eur J Pharmacol 298, 137–144 (1996).

54. M. R. Budhoo, J. M. Kellum, E. Woltering, S. Ashley, B. Bass, The 5-HT4 receptor mediates 5-hydroxytryptamine-induced rise in short circuit current in the human jejunum in vitro. Surgery 116, 396–400 (1994).

55. A. W. Baird, M. M. Skelly, D. P. O’Donoghue, K. E. Barrett, S. J. Keely, Bradykinin regulates human colonic ion transport in vitro. Br J Pharmacol 155, 558–566 (2008).

56. D. C. Manning, S. H. Snyder, J. F. Kachur, R. J. Miller, M. Field, Bradykinin receptor-mediated chloride secretion in intestinal function. Nature 299, 256–259 (1982).

57. Y. Wang, H. J. Cooke, R. Fertel, H. Su, Histamine augments colonic secretion in guinea pig distal colon. American journal of physiology 258, G432–G439 (1990).

58. G. Schultheiss, B. Hennig, W. Schunack, G. Prinz, M. Diener, Histamine-induced ion secretion across rat distal colon: Involvement of histamine H1 and H2 receptors. Eur J Pharmacol 546, 161–170 (2006).

59. S. Yarrow, J. A. Ferrar, H. M. Cox, The effects of capsaicin upon electrogenic ion transport in rat descending colon. Naunyn-Schmiedeberg’s Arch Pharmacol 344, 557– 563 (1991).

60. E. Weber, M. Neunlist, M. Schemann, T. Frieling, Neural components of distension-evoked secretory responses in the guinea-pig distal colon. Journal of Physiology 536, 741–751 (2001).

61. K. B. Seamon, W. Padgett, J. W. Daly, Forskolin: Unique diterpene activator of adenylate cyclase in membranes and in intact cells. Proc Natl Acad Sci U S A 78, 3363– 7 (1981).

62. R. J. Bridges, W. Rummel, B. Simon, Forskolin induced chloride secretion across the isolated mucosa of rat colon descendens. Naunyn Schmiedebergs Arch Pharmacol 323, 355–60 (1983).

63. L. Wang, A. M. Stanisz, B. K. Wershil, S. J. Galli, M. H. Perdue, Substance P induces ion secretion in mouse small intestine through effects on enteric nerves and mast cells. Am J Physiol Gastrointest Liver Physiol 269, G85–92 (1995).

64. M. Riegler, I. Castagliuolo, P. T. C. So, M. Lotz, C. Wang, M. Wlk, T. Sogukoglu, E. Cosentini, G. Bischof, G. Hamilton, B. Teleky, E. Wenzl, J. B. Matthews, C. Pothoulakis, Effects of substance P on human colonic mucosa in vitro. Am J Physiol Gastrointest Liver Physiol 276, G1473–83 (1999).

65. T. J. Park, C. Comer, A. Carol, Y. Lu, H. S. Hong, F. L. Rice, Somatosensory organization and behavior in naked mole-rats: II. Peripheral structures, innervation, and selective lack of neuropeptides associated with thermoregulation and pain. Journal of Comparative Neurology 465, 104–120 (2003).

66. R. Buffenstein, V. Amoroso, B. Andziak, S. Avdieiev, J. Azpurua, A. J. Barker, N. C. Bennett, M. A. Brieño-Enríquez, G. N. Bronner, C. Coen, M. A. Delaney, C. M. Dengler-Crish, Y. H. Edrey, C. G. Faulkes, D. Frankel, G. Friedlander, P. A. Gibney, V. Gorbunova, C. Hine, M. M. Holmes, J. U. M. Jarvis, Y. Kawamura, N. Kutsukake, C. Kenyon, W. T. Khaled, T. Kikusui, J. Kissil, S. Lagestee, J. Larson, A. Lauer, L. A. Lavrenchenko, A. Lee, J. B. Levitt, G. R. Lewin, K. N. Lewis Hardell, T. H. D. Lin, M. J. Mason, D. McCloskey, M. McMahon, K. Miura, K. Mogi, V. Narayan, T. P. O’Connor, K. Okanoya, M. J. O’Riain, T. J. Park, N. J. Place, K. Podshivalova, M. E. Pamenter, S. J. Pyott, J. Reznick, J. G. Ruby, A. B. Salmon, J. Santos-Sacchi, D. K. Sarko, A. Seluanov, A. Shepard, M. Smith, K. B. Storey, X. Tian, E. N. Vice, M. Viltard, A. Watarai, E. Wywial, M. Yamakawa, E. D. Zemlemerova, M. Zions, E. S. J. Smith, The naked truth: a comprehensive clarification and classification of current ‘myths’ in naked mole-rat biology. Biological Reviews 97, 115–140 (2022).

67. G. E. Holle, Changes in the structure and regeneration mode of the rat small intestinal mucosa following benzalkonium chloride treatment. Gastroenterology 101, 1264–73 (1991).

68. N. Miron, V. Cristea, Enterocytes: Active cells in tolerance to food and microbial antigens in the gut. [Preprint] (2012). 10.1111/j.1365-2249.2011.04523.x.

69. A. Bosi, D. Banfi, M. Bistoletti, L. M. Catizzone, A. M. Chiaravalli, P. Moretto, E. Moro, E. Karousou, M. Viola, M. C. Giron, F. Crema, C. Rossetti, G. Binelli, A. Passi, D. Vigetti, C. Giaroni, A. Baj, Hyaluronan Regulates Neuronal and Immune Function in the Rat Small Intestine and Colonic Microbiota after Ischemic/Reperfusion Injury. Cells 11, 3370 (2022).

70. M. Mitjans, R. Ferrer, Morphometric Study of the Guinea Pig Small Intestine during Development. Microsc Res Tech 63, 206–214 (2004).

71. J. Zhao, D. Liao, J. Yang, H. Gregersen, Biomechanical remodelling of obstructed guinea pig jejunum. J Biomech 43, 1322–1329 (2010).

72. R. Moore, C. Pothoulakis, J. Thomas Lamont, S. Carlson, J. L. Madara, J. L. Madara C difficile, C. difficile toxin A increases intestinal permeability and induces Cl-secretion. Am J Physiol 259, G165–72 (1990).

73. L. Feighery, A. Smyth, S. Keely, A. W. Baird, W. T. O’Connor, J. J. Callanan, D. J. Brayden, Increased intestinal permeability in rats subjected to traumatic frontal lobe percussion brain injury. Journal of Trauma - Injury, Infection and Critical Care 64, 131– 137 (2008).

74. B. I. Polentarutti, A. L. Peterson, À. K. Sjöberg, E. K. I. Anderberg, L. M. Utter, A. L. B. Ungell, Evaluation of viability of excised rat intestinal segments in the Ussing chamber: Investigation of morphology, electrical parameters, and permeability characteristics. Pharm Res 16, 446–54 (1999).

75. A. A. Livanova, A. A. Fedorova, A. V. Zavirsky, I. I. Krivoi, A. G. Markov, Dose- and Segment-Dependent Disturbance of Rat Gut by Ionizing Radiation: Impact of Tight Junction Proteins. Int J Mol Sci 24, 1753 (2023).

76. R. M. Van Elburg, J. J. Uil, C. J. J. Mulder, H. S. A. Heymans, Intestinal permeability in patients with coeliac disease and relatives of patients with coeliac disease. Gut 34, 354–7 (1993).

77. A. M. González-Castro, C. Martínez, E. Salvo-Romero, M. Fortea, C. Pardo-Camacho, T. Pérez-Berezo, C. Alonso-Cotoner, J. Santos, M. Vicario, Mucosal pathobiology and molecular signature of epithelial barrier dysfunction in the small intestine in irritable bowel syndrome. [Preprint] (2017). 10.1111/jgh.13417.

78. A. W. Baird, M. M. Skelly, D. P. O’Donoghue, K. E. Barrett, S. J. Keely, Bradykinin regulates human colonic ion transport in vitro. Br J Pharmacol 155, 558–66 (2008).

79. J. Yi, Z. Bertels, J. S. Del Rosario, A. J. Widman, R. A. Slivicki, M. Payne, H. M. Susser, B. A. Copits, R. W. Gereau, Bradykinin receptor expression and bradykinin-mediated sensitization of human sensory neurons. Pain 165, 202–215 (2024).

80. Y. Kawamura, K. Oka, T. Semba, M. Takamori, Y. Sugiura, R. Yamasaki, Y. Suzuki, T. Chujo, M. Nagase, Y. Oiwa, S. Fujioka, S. Homma, Y. Yamamura, S. Miyawaki, M. Narita, T. Fukuda, Y. Sakai, T. Ishimoto, K. Tomizawa, M. Suematsu, T. Yamamoto, H. Bono, H. Okano, K. Miura, Cellular senescence induction leads to progressive cell death via the INK4a-RB pathway in naked mole-rats. EMBO J 42, e111133 (2023).

81. K. N. Lewis, N. D. Rubinstein, R. Buffenstein, A window into extreme longevity; the circulating metabolomic signature of the naked mole-rat, a mammal that shows negligible senescence. Geroscience 40, 105–121 (2018).

82. M. B. Hansen, A. B. Witte, The role of serotonin in intestinal luminal sensing and secretion. [Preprint] (2008). 10.1111/j.1748-1716.2008.01870.x.

83. T. J. Park, Y. Lu, R. Jüttner, E. S. J. Smith, J. Hu, A. Brand, C. Wetzel, N. Milenkovic, B. Erdmann, P. A. Heppenstall, C. E. Laurito, S. P. Wilson, G. R. Lewin, Selective inflammatory pain insensitivity in the African naked mole-rat (Heterocephalus glaber). PLoS Biol 6, 0156–0170 (2008).

84. E. S. J. Smith, G. R. C. Blass, G. R. Lewin, T. J. Park, Absence of histamine-induced itch in the African naked mole-rat and “rescue” by Substance P. Mol Pain 6, 29 (2010).

85. D. I. Macdonald, M. Jayabalan, J. Seaman, A. Nickolls, A. Chesler, Pain persists in mice lacking both Substance P and CGRPα signaling. Elife 13, RP93754 (2024).

86. A. N. Anbazhagan, S. Priyamvada, W. A. Alrefai, P. K. Dudeja, Pathophysiology of IBD associated diarrhea. Tissue Barriers 6, e1463897 (2018).

87. W. D. Chey, J. Kurlander, S. Eswaran, Irritable bowel syndrome: A clinical review. American Medical Association [Preprint] (2015). 10.1001/jama.2015.0954.

88. J. D. Wood, Neuropathophysiology of the Irritable Bowel Syndrome. J Clin Gastroenterol 35, S11–22 (2002).

89. K. F. Ostrom, J. E. Lavigne, T. F. Brust, R. Seifert, C. W. Dessauer, V. J. Watts, R. S. Ostrom, Physiological roles of mammalian transmembrane adenylyl cyclase isoforms. [Preprint] (2022). 10.1152/physrev.00013.2021.

90. S. Pierre, T. Eschenhagen, G. Geisslinger, K. Scholich, Capturing adenylyl cyclases as potential drug targets. [Preprint] (2009). 10.1038/nrd2827.

91. T. V. Strong, K. Boehm, F. S. Collins, Localization of cystic fibrosis transmembrane conductance regulator mRNA in the human gastrointestinal tract by in situ hybridization. Journal of Clinical Investigation 93, 319–27 (1994).

92. R. A. Frizzell, J. W. Hanrahan, Physiology of epithelial chloride and fluid secretion. Cold Spring Harb Perspect Med 2, a009563 (2012).

93. A. Thomas, Y. Ramananda, K. S. Mun, A. P. Naren, K. Arora, AC6 is the major adenylate cyclase forming a diarrheagenic protein complex with cystic fibrosis transmembrane conductance regulator in cholera. [Preprint] (2018). 10.1074/jbc.RA118.003378.

94. X. Wang, K. Wang, X. Wu, W. Huang, L. Yang, Role of the cAMP–PKA–CREB–BDNF pathway in abnormal behaviours of serotonin transporter knockout mice. Behavioural Brain Research 419, 113681 (2022).

95. B. S. Dixon, Cyclic AMP selectively enhances bradykinin receptor synthesis and expression in cultured arterial smooth muscle: Inhibition of angiotensin II and vasopressin response. Journal of Clinical Investigation 93, 2535–44 (1994).

96. G. Bhave, W. Zhu, H. Wang, D. J. Brasier, G. S. Oxford, R. W. Gereau IV, cAMP-dependent protein kinase regulates desensitization of the capsaicin receptor (VR1) by direct phosphorylation. Neuron 35, 721–31 (2002).

97. N. Thevaranjan, A. Puchta, C. Schulz, A. Naidoo, J. C. Szamosi, C. P. Verschoor, D. Loukov, L. P. Schenck, J. Jury, K. P. Foley, J. D. Schertzer, M. J. Larché, D. J. Davidson, E. F. Verdú, M. G. Surette, D. M. E. Bowdish, Age-Associated Microbial Dysbiosis Promotes Intestinal Permeability, Systemic Inflammation, and Macrophage Dysfunction. Cell Host Microbe 21, 570 (2018).

98. Z. Zhou, C. Bian, Z. Luo, C. Guille, E. Ogunrinde, J. Wu, M. Zhao, S. Fitting, D. L. Kamen, J. C. Oates, G. Gilkeson, W. Jiang, Progesterone decreases gut permeability through upregulating occludin expression in primary human gut tissues and Caco-2 cells. Sci Rep 9, 8367 (2019).

99. X. Yang, Y. Guo, J. He, F. Zhang, X. Sun, S. Yang, H. Dong, Estrogen and estrogen receptors in the modulation of gastrointestinal epithelial secretion. Oncotarget 8, 97683–97692 (2017).

100. V. M. Kovalzon, O. A. Averina, V. A. Minkov, A. A. Petrin, M. Yu. Vysokikh, Unusual Correlation between Rest–Activity and Body Temperature Rhythms in the Naked Mole Rat (Heterocephalus glaber) as Compared to Five Other Mammalian Species. J Evol Biochem Physiol 56, 451–458 (2020).

101. G. P. Lambert, C. V. Gisolfi, D. J. Berg, P. L. Moseley, L. W. Oberley, K. C. Kregel, Selected contribution: Hyperthermia-induced intestinal permeability and the role of oxidative and nitrosative stress. J Appl Physiol 92, 1750–61 (2002).

102. K. Dokladny, P. L. Moseley, T. Y. Ma, Physiologically relevant increase in temperature causes an increase in intestinal epithelial tight junction permeability. Am J Physiol Gastrointest Liver Physiol 290, G204–12 (2006).

103. O. Inczefi, P. Bacsur, T. Resál, C. Keresztes, T. Molnár, The Influence of Nutrition on Intestinal Permeability and the Microbiome in Health and Disease. [Preprint] (2022). 10.3389/fnut.2022.718710.

104. A. C. Engevik, M. A. Engevik, Exploring the impact of intestinal ion transport on the gut microbiota. Comput Struct Biotechnol J 19, 134–144 (2021).

